# Comparative sugar utilisation and metabolism of mannose as co-substrate indicate flexibility in carbon metabolism in anaerobic gut fungi

**DOI:** 10.64898/2026.08.15.745028

**Authors:** Jessica L. Matthews, Hannah Haupt, Stephen C. Fry, Jolanda M. van Munster

## Abstract

Anaerobic gut fungi (AGF) are key degraders of plant biomass in ruminants, yet there is limited knowledge of how AGF respond to mixtures of plant-derived sugars. Here, we assessed monosaccharide and disaccharide utilisation by Neocallimastix frontalis CoB3, Caecomyces communis SHB, and Piromyces edwardsiae SHC, which are abundant in the rumen microbiome. While all AGF isolates shared a core set of sugars that supported growth, they had different hierarchies of uptake. Co-substrate experiments using glucose and lignocellulose-derived sugars revealed species-specific responses, with N. frontalis displaying a novel concentration-dependent co-utilisation of glucose and mannose, whereas growth of P. edwardsiae was inhibited under the same conditions, and C. communis exhibited growth inhibition in glucose and xylose co-substrate cultures. Together, these findings demonstrate functional diversity in monosaccharide and disaccharide metabolism amongst the AGF investigated here. Understanding such sugar utilisation phenotypes provides a foundation for evaluating AGF isolate suitability for lignocellulosic biomass valorisation.

## Introduction

Anaerobic gut fungi (AGF), phylum Neocallimastigomycota, are found in the digestive tracts of large variety of herbivores (Hanafy *et al.,* 2020). They were first isolated from the rumen (Braune, 1913; Orpin, 1975), where they are members of a complex microbial community that converts plant materials into nutrients the host animal can absorb, such as short-chain fatty acids (SCFAs) and microbial peptides (Tamminga and Van Vuuren, 1988). In the rumen, AGF are the primary colonisers of the ingested plant material (Grenet and Barry, 1988; Akin and Borneman, 1990) and are central to its degradation by physical and enzymatic action (Bauchop, 1979; Seppälä *et al.,* 2017).

The host diet dictates substrate availability for the rumen microbial community to consume, influencing the composition of the microbial community (Han *et al.,* 2019; Jones *et al.,* 2023). In the early life of the host animal when the diet is solely milk, AGF are borderline undetectable (Jones *et al.,* 2023). Once the host animal is weaned and the rumen and its microbial community are established, AGF account for approximately 10 % of the total microbial biomass when the host animal diet is forage-based (Nagaraja, 2016). However, when the host animal diet is supplemented with starch, AGF abundance decreases (Han *et al.,* 2019). On a genus-level, *Neocallimastix*, *Piromyces,* and *Orpinomyces* are seemingly more resistant to starch supplementation, with their relative abundance increasing when starch is added to their hosts’ diet (Hess *et al.,* 2020). Current knowledge of AGF genus abundance in host animals is largely limited to metagenomic analyses (Meili *et al.,* 2024); therefore, it is unclear whether the observed difference in AGF genera composition on different diets is due to substrate utilisation capacity, tolerance to changing pH, or a shift in microbial partnerships (Fliegerova *et al.,* 2021).

AGF have garnered great interest for their fibre degradative capacity, as despite their relatively small proportion of the microbiome compared to bacteria (∼50 %) (Nagaraja, 2016), AGF elimination *in vitro* and *in vivo* has been shown to significantly reduce fibre degradation (Theodorou *et al.,* 1996) - indicating they are key to this process. With the advancement of molecular tools in recent years, genomic and transcriptomic studies have revealed that AGF possess and express a large and diverse repertoire of carbohydrate-degrading enzymes (Youssef *et al.,* 2013; Solomon *et al.,* 2016; Haitjema *et al.,* 2017), further increasing interest in both their role in the rumen microbiome and their potential applications in biorefinery (Henske *et al.,* 2018b). Consequently, many studies of AGF have focused on their degradation of and growth on complex biomass (Gruninger *et al.,* 2018; Lillington *et al.,* 2021; Lankiewicz *et al.,* 2023). However, AGF primary metabolism and substrate preferences are poorly understood, especially the recognition, uptake and metabolism of monosaccharides and disaccharides in complex environments.

Here, the innate ability of three AGF isolates to utilise sugars that may be expected to be available in the rumen environment was investigated, focussing on AGF that are abundantly present in agriculturally important ruminants, and belonging to three different genera: *Neocallimastix frontalis* CoB3, *Caecomyces communis* SHB, and *Piromyces edwardsiae* SHC. From this, variable growth and uptake strategies of AGF isolates in a heterogeneous carbon source environment with mixtures of metabolisable and non-metabolisable sugars were identified. This included strong responses of all three fungi to some of the monosaccharides that they were unable to grow on as a single carbon source, such as the uptake and metabolism of mannose as a co-substrate. Furthermore, it was demonstrated that the substrate complexity of the inoculum has limited influence on the substrate utilization of subsequent cultures, in contrast to what was previously demonstrated in the rumen bacterium *Bacteroides* (Klassen *et al.,* 2021).

Together, these results provide an important phenotypic baseline to understand growth and primary metabolism of these AGF in ecological environments that contain complex mixtures of carbohydrate substrates, such as the rumen of herbivores.

## Materials and Methods

### Fungal cultivation

Axenic fungal isolates of *N. frontalis* CoB3, *C. communis* SHB and *P. edwardsiae* SHC (Shen *et al.,* 2026) were cultured under anaerobic conditions in medium C, modified from Orpin’s formulation to replace BactoCasitone with tryptone (Theodorou, Brookman, and Trinci, 2005). Clarified rumen fluid was obtained from a herd of rumen-cannulated Jersey cows on a seasonal diet (Glasgow University Farm). A colorimetric redox indicator, rezasurin (1 g L^−1^), was added to the medium to confirm anaerobic status.

Isolates were maintained by bi-weekly passage of 5 % fungal culture (v/v) into fresh medium C, with 1 % (w/v) milled wheat straw (0.5 mm particle size) as the carbon source. To prevent bacterial contamination, chloramphenicol was added to cultures to 50 µg mL^−1^. Fungal cultures were grown at 39 °C with stationary incubation.

### Measuring fungal growth

The production of fermentation gases is used as the proxy for measuring anaerobic fungal growth (Theodorou *et al.,* 1995; Wilken *et al.,* 2020). Fermentation gas pressure was measured every 24 hours via a Vernier LabQuest 2 data logger equipped with a pressure sensor. Before gas pressure was measured, cultures were transferred to a 39 °C water bath, and the Hungate tube caps and septa were ethanol sterilized. After the measurement was recorded, the fermentation gases were vented to prevent their accumulation from inhibiting fungal growth (Joblin and Naylor, 1993). Gas accumulation was measured every 24 hours.

The pressure was normalized using an abiotic ‘blank’ medium as a control. After the final gas pressure measurement was recorded, the Hungate tubes were opened, and the culture pH was measured immediately, using an Orionstar A111 benchtop pH meter equipped with a semi-micro pH probe (Fisherbrand FB68801).

### Monitoring fungal uptake of sugars

For time-course sampling, 0.1-mL aliquots of medium were removed from fungal and blank cultures every 24 hours after gas pressure was recorded. For end-point sampling, a 2-mL aliquot of medium was removed before pH measurement. These aliquots were immediately placed on ice to arrest sugar uptake by fungal cells also present in the culture medium sample. Samples were stored at –20 °C until use, when they were thawed at room temperature and centrifuged briefly at 11,000 *g* to pellet any fungal biomass present.

To separate and detect sugars remaining in the culture medium, 3 µL of an aliquot of culture medium and 2.5 µL of a sugar marker mixture (each sugar present corresponding to the experimental condition, at 0.1 % (w/v)) were dried onto a Merck silica-gel 60 TLC plate. The TLC plate was developed in ethyl acetate/pyridine/acetic acid/water (6:3:1:1 by volume), using two ascents. The plate was stained with thymol/H_2_SO_4_/ethanol (0.5:5:95, w/v/v) and heated at 105 °C for 15 minutes (Franková & Fry, 2021).

### Growth assay of AGF isolates on monosaccharides and disaccharides

A selection of constituent monosaccharides and disaccharides of lignocellulose (D-glucose, D-xylose, L-arabinose, D-galactose, D-mannose, L-rhamnose, D-glucuronic acid, D-galacturonic acid, cellobiose) and those commonly found in the rumen environment (maltose, sucrose, and lactose) were prepared by filter-sterilisation and added to medium C to a final concentration of 5 g L^−1^. Concentrated sugar stock solutions were stored at −20 °C.

These experimental media were inoculated with 5 % (v/v) 3-day-old fungal cultures grown on either wheat straw (0.5 mm particle size) or glucose (5 g L^−1^) prepared as above.

### Preferential uptake of metabolisable sugars

Sugar mixtures, namely “M1” (glucose, fructose, cellobiose, lactose) and “M2” (glucose, fructose, cellobiose, maltose, and lactose) were prepared with equal concentrations (w/w) of each component sugar. Each mixture was supplemented into the culture medium to a final concentration of 5 g L^−1^ (each sugar 1.25 g L^−1^ in “M1” and 1.0 g L^−1^ in “M2”) and inoculated with 3-day-old fungal cultures grown on wheat straw (0.5 mm particle size). The culture medium was sampled every 24 hours as described above.

### Recognition and response to ‘non-metabolisable’ sugars in the presence of glucose

All sugar stocks were prepared by filter-sterilisation. Glucose was supplemented to the medium C as the growth substrate to a final concentration of 5 g L^−1^. An additional sugar, either galactose, mannose, arabinose or xylose, was added to the culture medium to a final concentration of 2.5 g L^−1^. Then, cultures were inoculated with 5 % (v/v) 3-day-old fungal cultures which had been grown on wheat straw (0.5 mm particle size). The culture medium was sampled after the final gas pressure measurement as described above.

### Assessment of mannose concentrations on use as co-substrate with glucose and fructose

Mannose was supplemented at 2.5 g L^−1^ or 5 g L^−1^ to medium C together with either 2.5 g L^−1^ or 5 g L^−1^ glucose or fructose. For growth controls, culture medium was also prepared with each sugar as the sole carbon source at concentrations used in this experiment. These cultures were inoculated with 5 % (v/v) 3-day-old *N. frontalis* CoB3 cultures that had been grown on wheat straw. The culture medium was sampled every 24 hours and aliquots analysed by TLC. For quantification of fungal biomass, fungal biomass was collected by centrifugation of cultures at 10 min for 4500 rpm, and subsequently freeze dried and weighted.

### Potential co-substrate utilisation mechanisms of N. frontalis CoB3

^14^C-Radiolabelled sugars were utilised to detect to a high sensitivity whether mannose or arabinose metabolism occurs by *N. frontalis* CoB3 in the presence of glucose. [2-^14^C]Glucose (specific activity 2.07 MBq µmol^−1^) was used as received from the supplier. The other ^14^C-radiolabelled sugars were prepared as follows: [1-^14^C]arabinose (Amersham International, specific activity 2.035 MBq µmol^−1^) and [1-^14^C]mannose (Amersham International, specific activity 2.183 MBq µmol^−1^) stocks were purified by descending paper chromatography (57 cm × 46 cm, Whatman filter paper #1). To aid in identification of the sugars of interest, 2.5 µg of glucose, mannose, arabinose, and xylose were loaded as external markers onto the paper. The paper chromatogram was then developed in a solvent system of ethyl acetate/pyridine/water (8:2:1, v/v). Once developed in the solvent system, the external marker lanes were cut and stained using aniline hydrogen-phthalate (Fry, 1988), revealing the positions of the radiolabelled sugars of interest. The respective regions of the paper for mannose and arabinose were cut, and the radiolabelled sugars were eluted by multiple washings with deionised water until the paper was returned to background levels of radiation, assessed by a Geiger counter. The collected eluates were dried under vacuum and redissolved in deionised water. The activity concentration for both mannose and arabinose was determined by the scintillation-counting of triplicate small aliquots in OptiPhase HiSafe 3 scintillant fluid (PerkinElmer, Inc.) (10:1, v/v, scintillant : sample).

To have a baseline of metabolism end products which could be detected and quantified, control cultures were prepared supplemented with non-radioactive glucose (5 g L^−1^) and [^14^C]glucose. For co-sugar cultures, non-radioactive glucose was added to medium C (5 g L^−1^), and the additional non-radioactive sugar (either mannose or arabinose) was also added (2.5 g L^−1^) with the corresponding radiolabelled sugar, [^14^C]mannose or [^14^C]arabinose. Each radiolabelled sugar was added in 50 µL using a Hamilton syringe, before fungal inoculation. As the concentration of the radiolabelled sugars were negligible compared to the concentrations of non-radioactive sugar, the overall sugar concentrations were not adjusted for the added radiolabelled sugars.

Cultures were inoculated with 5 % (v/v) 3-day-old fungal cultures grown on wheat straw (0.5 mm particle size), and their gases were vented every 24 hours. Cultures were harvested at 120 hours as complete glucose removal from the culture medium was expected, by filtration through glass wool to separate the fungal biomass from the culture medium. For each biological replicate, duplicates of 0.5 mL culture medium filtrate were diluted with 1.5 mL of deionised water, to sufficiently dilute the oxygenated culture medium to prevent colour quenching during scintillation-counting. Then, the glass wool and fungal biomass was washed three times with 10 mL of 1X phosphate buffer solution (PBS) to remove all culture medium, with the three collections kept separate. 2 mL of each washing collection was also sampled in duplicate. Both the culture medium filtrate and washing samples were prepared for liquid scintillation counting as described above. 5 mL of the same scintillant fluid was added to the glass wool containing the fungal biomass, which was mixed thoroughly. Blanks of culture medium filtrate, 1X PBS, and glass wool were also prepared in triplicate.

The major metabolic products of AGF expected to be present in the culture medium filtrate were lactate, ethanol, succinate, and short chain fatty acids (SCFAs) (acetate and formate) (Wilken *et al.,* 2021). To identify if the radiolabelled sugar was metabolised into any of these products, the culture medium filtrate was sub-aliquoted and prepared as described below:

Preparation 1) 100 µL of filtrate was added to 400 µL deionised water to scintillation-count the radioactivity present in whole filtrate (non-volatile, neutral volatile, and acidic volatile solutes).

Preparation 2) 100 µL of filtrate was added to 100 µL 50 % acetic acid and then dried. This ensured protonation of the organic acids to allow their removal by evaporation, and selective recovery and counting of non-volatile solutes.

Preparation 3) 100 µL of filtrate was added to 100 µL 1 M sodium hydroxide and dried to remove neutral volatiles (e.g. ethanol) while retaining acidic solutes in their non-volatile salt form. To allow for proper scintillation-counting and quantification, these salts were then re-acidified with 100 µL 50 % acetic acid to regenerate the acids, then 400 µL of deionised water was added. This preparation contained both non-volatile solutes (e.g. sugars and lactate) and SCFAs (e.g. acetate and formate).

Each sample was prepared in triplicate for each harvested culture and prepared for liquid scintillation counting as previously described. The radioactivity counts of the preparations were then used to calculate the broad biochemical nature of ^14^C-labelled solutes as described in Table 1.

**Table 1:** Description of calculations to identify and quantify the biochemical nature of ^14^C metabolites in the filtrate collected from N. frontalis CoB3 cultures.

| Biochemical nature | Calculation (blank corrected cpm) |
| --- | --- |
| Non-volatile (e.g. lactate, succinate and sugar) | Preparation 2 |
| Neutral volatile (e.g. ethanol) | [Preparation 1] - [Preparation 3] |
| Acidic volatile (e.g. acetate and formate) | [Preparation 3] - [Preparation 2] |

High-voltage paper electrophoresis was used to further investigate if the ^14^C recovered in the non-volatile fraction consisted of anionic metabolites e.g. lactate and/or the dosed ^14^C-sugar. 300 µL filtrate was concentrated under vacuum and redissolved in 45 µL of 0.5 % chlorobutanol. External markers were glucose (10 µg) plus a mixture of acids: succinic, citric, L-tartaric, malonic, pyruvic (each 0.5 µmol) and lactic (1 µmol). The whole 45 µL of each sample was loaded onto Whatman paper #3 (52 cm × 46 cm) alongside the above external marker mixture. To visualise the migration of the loadings down the paper, 2 µL of internal marker (0.4 µg of orange G and 0.1 µg of xylene cyanol) was also added to each sample and marker. Then, the paper was wetted with pH 3.5, acetic acid/pyridine/water electrophoresis buffer (10:1:89, v/v) and placed in a tank containing immiscible coolant of white spirit. This paper was electrophoresed at 2.5 kV for 55 minutes (Fry, 2020). The paper was dried, and the external markers were cut and stained firstly with bromophenol blue to reveal the anions. To confirm localisation of neutral sugars near the origin, these markers were then stained with silver nitrate (Fry, 2020), which also stains some of the anionic markers.

The unstained electrophoretogram sample lanes were cut into 3 cm x 2 cm sections each of which was added to 2 mL scintillation fluid for liquid scintillation counting; blanks of clean Whatman paper #3 were prepared in triplicate. From the results of the counting, the becquerels per centimetre of each sample lane was calculated to localise and quantify the peaks of radioactivity and identify their position relative to the components of interest from the external and internal markers (Supplementary Figure 4).

### Statistical Analysis

Per dataset, each AGF isolate’s respective results were assessed independently for normality using QQ-plot and Shapiro-Wilk. If the normality of residuals correlated linearly and the Shapiro-Wilk test was not significant (p-value > 0.05), parametric statistical tests were performed: either Student’s t-test, one-way or two ANOVA (significant p-value ≤ 0.05). To assess where the significant differences between means lie if the ANOVA was significant (p-value ≤ 0.05), Tukey HSD was performed. If the data were not normally distributed and the Shapiro-Wilk test was significant (p-value ≤ 0.05), non-parametric tests were performed: Mann-Whitney U test or the Kruskal-Wallis test followed by pairwise Wilcoxon test if significant (significant p-value ≤ 0.05).

## Results

### Growth assay of AGF isolates on monosaccharides and disaccharides

The growth of AGF was assessed on sugars likely to be available in the rumen environment from the feed intake of the host. As AGF are most abundant in the rumen when the host animal diet is forage-based, most of the screened sugars were plant-derived: glucose, cellobiose, fructose, sucrose, xylose, arabinose, galactose, mannose, rhamnose, glucuronic acid, and galacturonic acid. Maltose was included as the disaccharide to represent a starch diet, and lactose as the available sugar in milk. To see if the culture substrate used to maintain AGF isolates influences the sugar utilisation of the subsequent cultures, we performed this screen using inoculum grown either on wheat straw or on glucose.

All AGF isolates screened here could utilise glucose, cellobiose, fructose, and lactose for growth. However, the growth performance of each respective isolate did vary between these sugars (Fig. 1A,B). *P. edwardsiae* isolate SHC exhibited minimal growth on fructose as although these cultures generated visible fungal biomass, they produced minimal fermentation gas and no significant change in pH was observed compared to the ‘blank’ control cultures (Fig. 1D).

**Figure 1.**
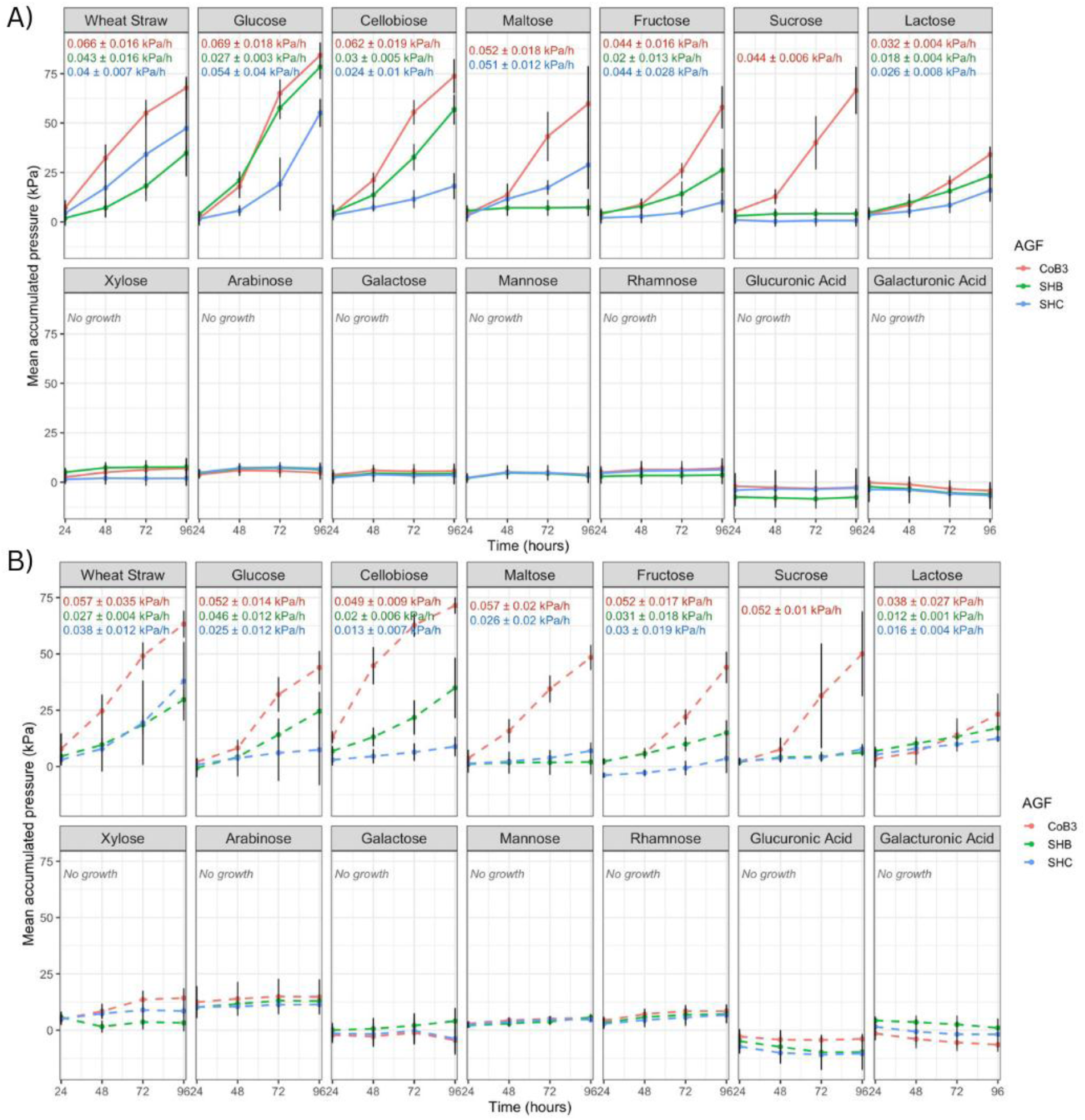
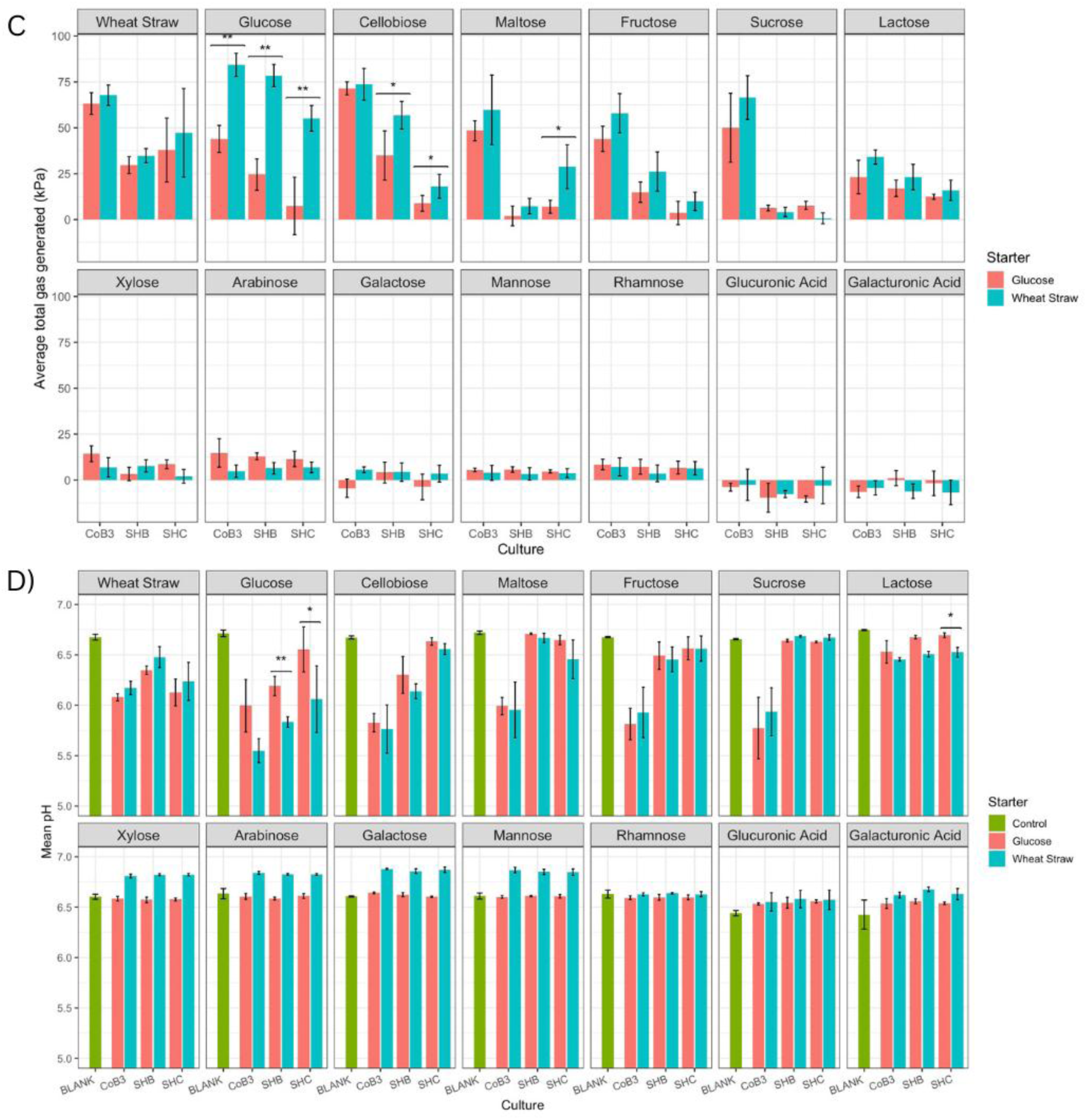
Growth of fungal isolates N. frontalis CoB3, C. communis SHB, and P. edwardsiae SHC on various soluble sugars. A) Fermentation gas accumulation by culture from inoculum with wheat straw as carbon source and B) inoculum with glucose as carbon source. Carbon sources used in the culture are indicated in panel headers, insets display exponential rate of growth for conditions in which fungal growth occurred. C) Total accumulation of fermentation gas and D) pH after 96 h after fungal inoculation; the control had no fungal inoculation. All values reported are mean ± stdev (n=6). Significance is reported for comparison of inoculation from wheat straw-pregrown versus glucose-pregrown cultures by Mann-Whitney U test, with p-values given as, *** p < 0.001; ** 0.001 ≤ p < 0.01; * 0.01 ≤ p < 0.05.

Furthermore, all isolates exhibited poor growth on lactose relative to the other substrates they were able to utilise. Both total accumulated gas production and the exponential growth rate (calculated as described in Fig. S1) were approximately half of those observed for the wheat straw control (Fig. 1A,C). As a disaccharide of galactose and glucose, lactose utilisation could be energetically less favourable compared to glucose-glucose disaccharides, as only glucose can be utilised for growth and none of these isolates can utilise galactose as the sole carbon source (Fig. 1A). *C. communis* SHB growth was limited to these four sugars (glucose, fructose, cellobiose, and lactose); however, SHC could also utilise maltose. *N. frontalis* CoB3, the most versatile isolate assayed here, could also utilise maltose and sucrose. SHB’s limited growth substrate profile and inability to utilise maltose is reflective of other characterised isolates of the *Caecomyces* genus (Henske *et al.,* 2017).

The substrates that AGF isolates could utilise for growth did not change whether the culture was inoculated from fungi grown on wheat straw (lignocellulose) (Fig. 1A) or glucose (a simple sugar) (Fig. 1B). However, there was an effect on the total accumulated gas pressure (Fig. 1C): on glucose (CoB3, SHB, SHC), cellobiose (SHB, SHC), or maltose (SHC), cultures pre-grown on glucose showed significantly reduced total accumulated gas compared to when pre-grown on wheat straw (Fig. 1). The growth substrate of the inoculum did not affect the pH recorded for CoB3 on these substrates, but for both SHB and SHC the pH was significantly decreased when grown on glucose and lactose. This could be linked to the reduced rate of exponential growth recorded in these conditions, suggesting a possible difference in metabolic profile.

Overall, these results indicate that the AGF investigated here grow on a core set of sugars, whereby *N. frontalis* CoB3 and *P. edwardsiae* SHC have a slightly broader utilisation profile when ‘simple sugars’ (i.e. monosaccharides and disaccharides) are given as the sole carbon source.

### Preferential uptake of metabolisable sugars

When AGF digest complex plant material, in either the rumen or a renewables-based biotechnology process, simple sugars will be released and present as a mixture. Therefore, we assessed the AGF isolates’ preferential uptake of their metabolisable sugars. AGF isolates were grown on two different mixtures of the sugars identified to support fungal growth as the sole carbon source. Mixture 1, “M1”, contained the sugars all isolates could utilise: glucose, fructose, cellobiose, and lactose. Mixture 2, “M2”, also in addition included maltose, which both CoB3 and SHC could utilise as the sole carbon source. Whilst SHB and other isolates of this genus cannot utilise maltose as the sole carbon source, the *C. communis var. churrovis* genome does encode amylases (Henske *et al.,* 2017); therefore, it was also grown in this mixture.

For each AGF isolate, there was no significant difference in total accumulated gas pressure and pH between the two sugar mixtures (Fig 2). Despite having a final concentration of 5 g L^−1^ of sugar in both mixtures, the growth performance on the mixtures was worse for each AGF isolate than on each sugar alone (Fig. 1), except for lactose (all isolates), fructose (SHB and SHC), and cellobiose (SHC).

**Figure 2.**
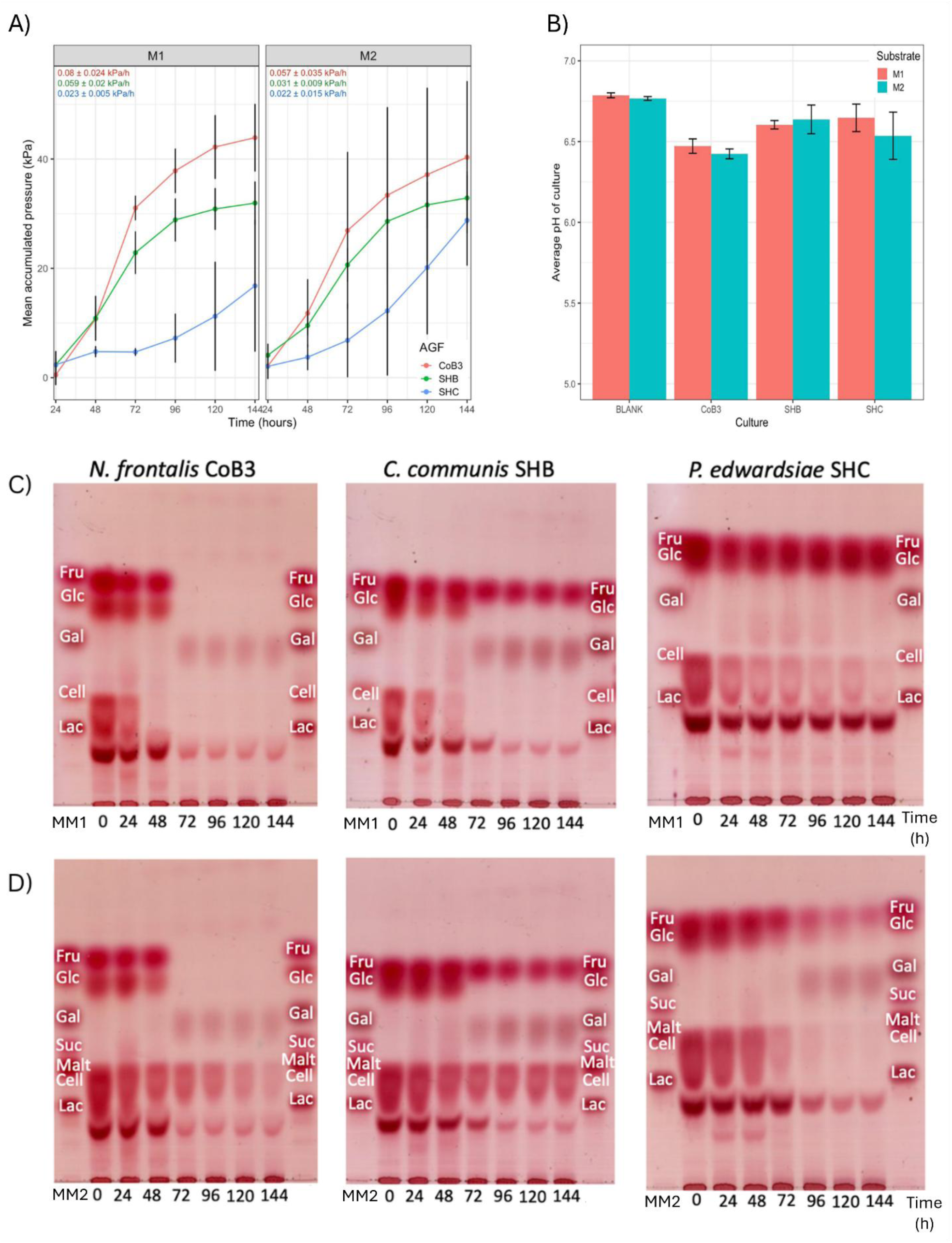
Growth of fungal isolates N. frontalis CoB3, C. communis SHB, and P. edwardsiae SHC on sugar mixtures “M1”: glucose, fructose, cellobiose, and lactose and “M2”: glucose, fructose, cellobiose, maltose, and lactose. The final sugar concentration of each sugar mixture was 5 g L^−1^. Values reported are mean ± stdev (n=5). A) Fermentation gas accumulation of cultures and the rate of exponential growth for each AGF isolate and B) pH of culture supernatant after 144 h of growth. The culture medium was sampled every 24 hours for analysis by thin-layer chromatography; a representative replicate of each fungal isolate N. frontalis CoB3, C. communis SHB, and P. edwardsiae SHC is shown grown on C) sugar mixture “M1” [marker mixture 1 (MM1) contains these sugars and galactose dissolved in water for reference] and D) and sugar mixture “M2” [marker mixture 2 (MM2) contains these sugars, galactose and sucrose dissolved in water for reference]. ‘0 h’ is a pooled sample of the culture medium supplemented with the sugar co-substrates before fungal inoculation.

To identify whether AGF isolates have preferential uptake of sugars and different strategies of sugar uptake (sequential vs. simultaneous), we removed aliquots of the culture medium every 24 hours for TLC analysis. This analysis revealed that the AGF isolates investigated here have different preferences and strategies for sugar uptake (Fig. 2C,D). Lactose was never completely removed from the culture medium in any condition tested, and its partial consumption was observed to be accompanied by the accumulation of galactose (Fig. 2C,D). This accumulation was expected, as the AGF isolates could not utilise galactose for growth (Fig. 1). The retention factors (*R*_F_s) of especially disaccharides in the complex sample matrix were slightly different from those in the standard; retention factors were confirmed independently using spike-ins (Fig. S2).

Grown on sugar mixture “M1”, *N. frontalis* CoB3 preferentially removed cellobiose first, removing it from the culture medium by 48 hours. Glucose and fructose were seemingly co-utilised, with both being completely removed from the medium by 72 hours. A reduction in lactose concentration was detected between 48 and 72 hours in both sugar mixtures.

Similarly, in “M2”, cellobiose was the first sugar CoB3 interacts with, by its hydrolysis into glucose. This is inferred from the observed accumulation of glucose at 24 hours compared to 0 hours (Fig. 2C,D), and using the position of cellobiose in “M1” on the same plate, as the separation of maltose and cellobiose is ambiguous in both water and rumen fluid, owing to their similar structures. The uptake of glucose was then preferred to fructose as observed by the relative intensity of the spot at 72 hours. Maltose was not completely removed from the culture medium at 144 hours, but more so than lactose (Fig. 2C,D).

*C. communis* SHB showed similar preferential uptake of sugar mixture “M1” to CoB3, but the uptake was slower, with the majority of cellobiose removed at 48 hours and complete removal from the culture medium occurring by 72 hours (Fig. 2C). Unlike CoB3, fructose remained present in the culture medium for the 144 hours measured. In sugar mixture “M2”, this preference for sugar uptake was retained (Fig. 2D). The additional maltose remained in the culture medium, as expected from the sole carbon source growth screen (Fig. 1), with fructose and some lactose.

Surprisingly, *P. edwardsiae* SHC only consumed cellobiose from sugar mixture “M1”. The majority of cellobiose disappeared within 24 hours; however, complete removal from the culture medium was not observed (Fig. 2C). The rest of the sugars (glucose, fructose, and lactose) were seemingly at consistent concentrations at each sampled time point, suggesting they were not removed from the culture medium. The lack of sugar uptake would explain the poor growth observed in the growth curve (Fig 2A). The addition of maltose to the sugar mixture seemingly rescued SHC growth (Fig 2A). From the TLC analysis, more sugar was removed from the culture medium; cellobiose, maltose, and glucose were mostly removed by 72 hours, and fructose consumption began at 72 hours, with lactose beginning 24 hours later (Fig. 2D). However, complete removal from the culture medium was not observed for any of these sugars.

### Recognition and response to ‘non-metabolisable’ sugars in the presence of glucose

As the primary colonisers of plant biomass (Grenet and Barry, 1988; Akin and Borneman, 1990), AGF will be in a highly diverse sugar environment of varying ratios of glucose, xylose, arabinose, mannose, and galactose (Sluiter *et al.,* 2010). However, the AGF isolate growth screen (Fig. 1) demonstrated that many of the lignocellulose-derived sugars cannot be utilised for growth when present as the sole carbon source. To assess how AGF growth is affected by the presence of sugars they cannot utilise as the sole carbon source, AGF growth was assayed in co-substrate cultures containing glucose (5 g L^−1^) supplemented with major lignocellulose-derived sugars (2.5 g L^−1^).

*N. frontalis* CoB3 was the only isolate assayed here to show clear evidence of co-substrate utilisation of glucose plus a different lignocellulose-derived sugar; in this culture, the presence of mannose significantly increased total gas accumulation and pH reduction compared to glucose-only cultures (Fig. 3A,B), indicating a stimulation of growth. TLC analysis (Fig. 3C) revealed that all the glucose and the majority of mannose were removed from the culture medium. Whilst CoB3 is not the first strain of *N. frontalis* reported to synergistically utilise mannose and glucose, *N. frontalis* CoB3 cannot utilise mannose as the sole carbon source (Fig. 1) unlike the reported *N. frontalis* strain RE1 (Stewart *et al.,* 1995). Therefore, *N. frontalis* CoB3 co-substrate dependency on glucose for mannose utilisation is a novel metabolic trait, and to our knowledge the first report of a Neocallimastigomycota isolate able to metabolise sugars only via a strict co-substrate dependency.

**Figure 3.**
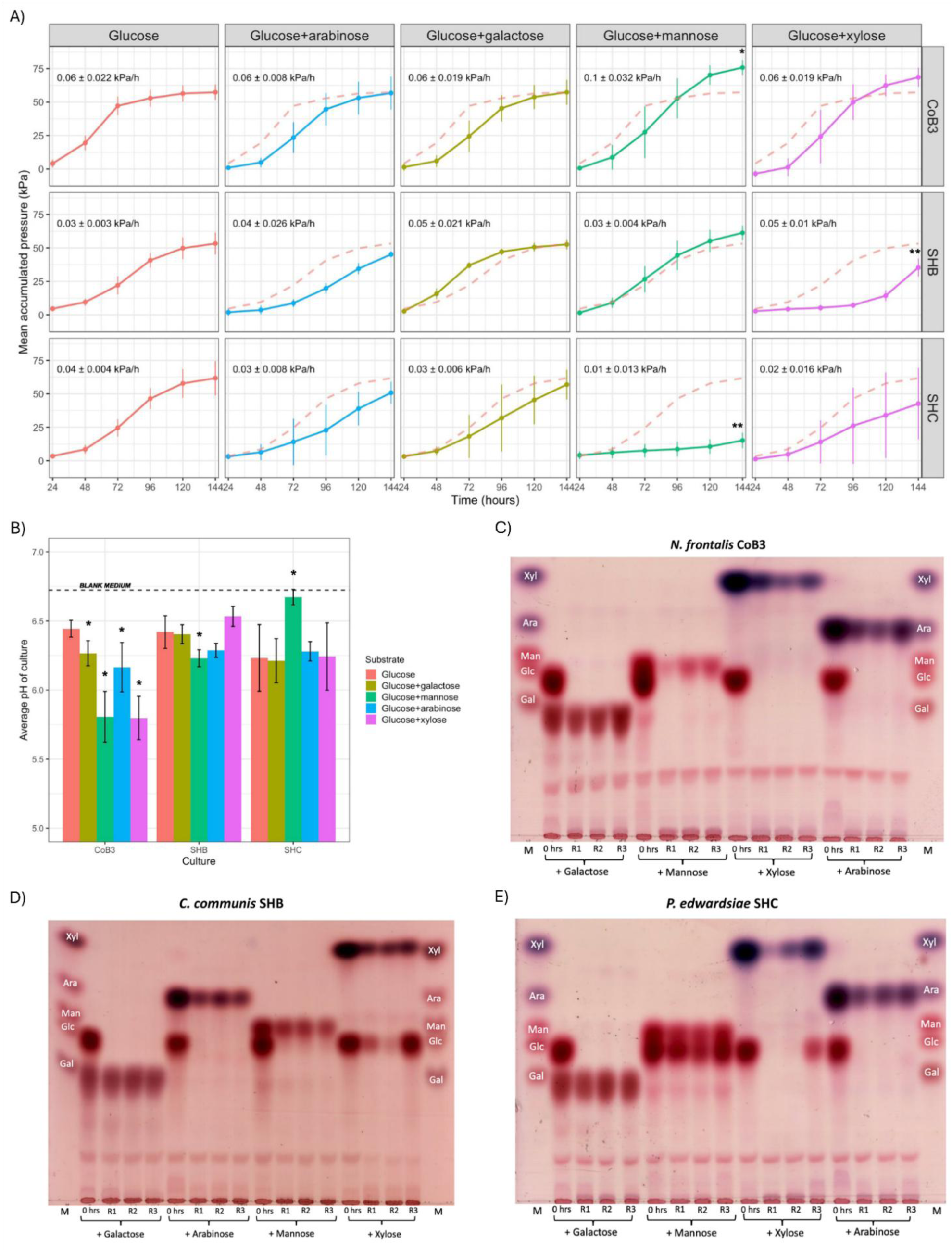
Growth of fungal isolates N. frontalis CoB3, C. communis SHB, and P. edwardsiae SHC on glucose + lignocellulose-derived sugars. Glucose was added to a final concentration of 5 g L^−1^ and the other lignocellulose-derived sugar (if any) to a final concentration of 2.5 g L^−1^. Values reported are mean ± stdev (n=5). A) Fermentation gas accumulation of cultures and rate of exponential growth; B) pH of culture supernatant 144 h after fungal inoculation. Significance values reported are for AGF isolates on glucose + other lignocellulose-derived sugar cultures versus glucose-only cultures; one-way ANOVA and Tukey HSD analysis was used to assess all AGF isolates’ gas pressure and the pH of SHB and SHC, and for CoB3 pH Kruskal-Wallis and pairwise Wilcoxon test was used. For all statistical tests p-values given as *** p < 0.001; ** 0.001 ≤ p < 0.01; * 0.01 ≤ p < 0.05. Three biological replicates with the least variation of total fermentation gas production were chosen for thin-layer chromatography analysis: C) N. frontalis CoB3 D) C. communis SHB E) P. edwardsiae SHC. “M” is a marker mixture of glucose, galactose, mannose, xylose, and arabinose dissolved in water. ‘0 h’ is a pooled sample of the culture medium supplemented with the sugar co-substrates before fungal inoculation. 144 h after fungal inoculation all replicates’ culture media were sampled.

Although *C. communis* SHB and *P. edwardsiae* SHC did not show evidence of co-substrate utilisation of another lignocellulose-derived sugar with glucose, their growth was hindered by the presence of some of these sugars (Fig. 3A,B). Conversely to *N. frontalis* CoB3, *P. edwardsiae* SHC growth was significantly reduced by the presence of mannose as shown in the reduction in total gas accumulation and culture pH compared to glucose-only (Fig. 3A,B). Furthermore, the TLC analysis showed limited removal of either glucose or mannose from the culture medium, suggesting that somehow the presence of mannose prevented SHC glucose uptake (Fig. 3E). The presence of xylose showed a similar impact on growth and glucose uptake for SHB (Fig. 3A,D); however, the reduction of glucose uptake was more visibly variable between biological replicates (Fig. 3D). Surprisingly, the TLC analysis revealed similar variable xylose uptake for also CoB3 and SHC between their biological replicates (Fig. 3D,E). Seemingly, xylose did not have a significant effect on the total fermentation gas production recorded for CoB3, but for SHC it did correlate with variable fermentation gas production (Fig. 3A). The presence of residual glucose in the culture medium and the reduced fungal growth suggests that xylose and mannose disrupted glucose sensing, uptake, or metabolism - for example by competing for glucose transport into fungal cells in SHB and SHC.

The addition of galactose or arabinose as a co-substrate did not impact the total fermentation gas production or the total removal of glucose from any of the AGF isolates here (Fig. 3). Visual inspection of AGF growth curves suggests, even for lignocellulose-derived sugars which did not significantly affect the total fermentation gas accumulation, glucose-only cultures reached exponential growth sooner than co-substrate cultures (Fig. 3A). There was no evidence of galactose uptake by any of the AGF isolates here from TLC analysis (Fig. 3C-E), but the data did suggest minimal arabinose removal from the culture medium, especially for CoB3 (Fig. 3C-E). This was investigated further by time course sampling CoB3 co-substrate cultures of glucose and arabinose, but this showed no clear evidence of arabinose uptake (Fig. S3).

In summary, AGF isolates *N. frontalis* CoB3, *C. communis* SHB, and *P. edwardsiae* SHC differ in their responses to previously assigned ‘non-metabolised’ lignocellulose-derived sugars when grown on glucose, which vary from repression of growth to a co-substrate dependent uptake and metabolic use of these ‘non-metabolised’ sugar. This suggests these AGF isolates have intricate underlying regulatory or metabolic networks enabling them to sense and respond to simple sugars, irrespective of their ability to utilise these sugars as the sole carbon source in an axenic culture.

### Assessment of the effect of glucose and mannose concentrations on N. frontalis CoB3 co-substrate utilisation

To investigate in further detail the synergistic utilisation of glucose and mannose observed in *N. frontalis* CoB3, a co-substrate screen at various concentrations and ratios of glucose to mannose (1:1, 2:1, and 1:2) was performed to assess if this co-substrate utilisation is concentration dependent. To ascertain if the co-substrate utilisation occurs simultaneously or sequentially, sugars in the culture medium were analysed every 24 hours.

Effective co-sugar utilisation of mannose in the presence of glucose *N. frontalis* CoB3 seemed to depend on the glucose concentration being equal or in excess of mannose, and the mannose concentration below 5 g L^−1^ (Fig. 4, Fig S4). In cultures with a mannose concentration of 5 g L^−1^ of mannose, growth was significantly delayed or decreased compared to glucose-only cultures based on fermentation gas and pH (Fig. 4A,B). TLC analysis revealed that at these concentration ratios, glucose remains in the culture medium suggesting that mannose diminishes glucose removal, corresponding to reduced CoB3 growth (Fig. 4C,E). However, when glucose was in excess to mannose, growth was significantly increased (Fig. 4A,B). Both sugars were progressively removed from the culture at each time point sampled (Fig. 4C), but glucose was preferentially removed first. The presence of 2.5 g L^−1^ mannose may have delayed glucose uptake by up to 24 hours compared to the glucose-only culture (5 g L^−1^) (Fig. 4C, D), but the transition to exponential growth was still observed at similar time points (Fig. 4A,B), further suggesting that mannose is metabolised by CoB3 as there is no hinderance of growth from the reduced glucose removal. Fungal biomass accumulation (Fig. S4B,C), assessed in replicate cultures containing glucose and mannose that were sacrificed at three different timepoints (Fig. S4A), followed trends observed for fermentation gas production (Fig. S4A) and pH (Fig. S4D). This confirmed that these effects were not only a result of relative changes to fungal metabolism, but a direct result of fungal growth.

**Figure 4.**
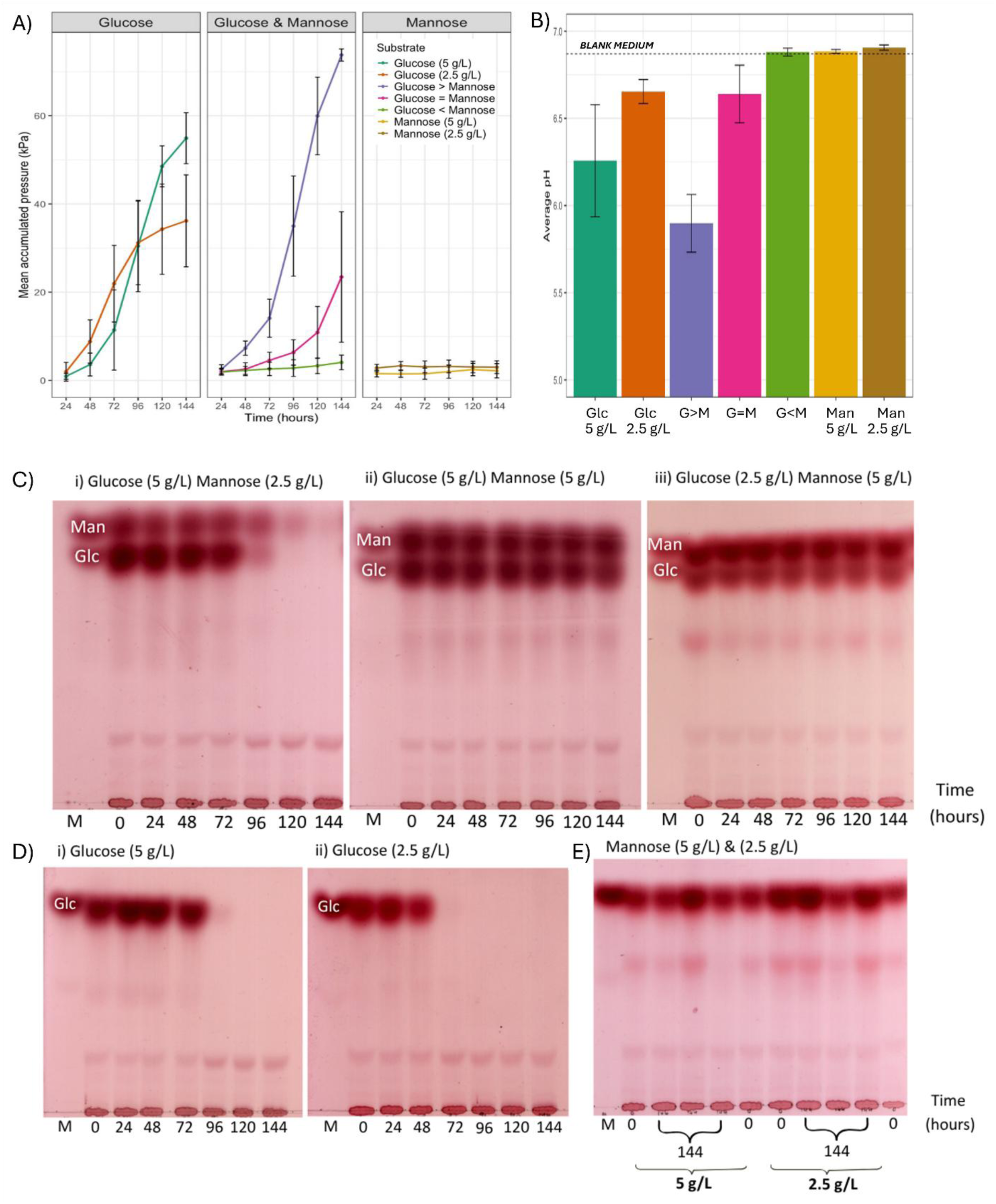
Growth of fungal isolate N. frontalis CoB3 in co-substrate cultures of glucose and mannose. The respective final concentrations of sugars in cultures were: glucose (5 g L^−1^) > mannose (2.5 g L^−1^), glucose = mannose (both 5 g L^−1^), and glucose (2.5 g L^−1^) < mannose (5 g L^−1^). Values reported are mean ± stdev (n=5). A) Fermentation gas accumulation of cultures; B) pH of culture supernatant 144 h after fungal inoculation. C) thin-layer chromatography of culture filtrate of a representative replicate culture. i) glucose > mannose, ii) glucose = mannose, iii) glucose < mannose, D) glucose control i) 5 g L^−1^ and ii) 2.5 g L^−1^. Cultures with mannose as the only carbon source did not show evidence of growth so only E) 144 h of three biological replicates was analysed. ‘0 h’ is a pooled sample of the culture medium supplemented with the sugar co-substrates before fungal inoculation. Abbreviations: Glc, Glucose: Man, Mannose.

While *N. frontalis* CoB3 grows well on fructose as sole carbon source (Fig. 1), cultures did not grow when containing a combination of fructose and mannose, regardless of ratios and concentrations varying from 2.5 g L^−1^ fructose with 5 g L^−1^ mannose to 5 g L^−1^ fructose with 2.5 g L^−1^ mannose (Fig. S5). Growth was monitored for an extended time, 28 days, during which no changes were observed, excluding a long lag phase precluding observation of growth. This indicates that use of mannose is not enabled by presence of a generic carbon source, but specifically depend on the presence of glucose. The observed growth repression in cultures with fructose and mannose suggests further complex metabolic interactions may take place.

In summary, this data strongly suggests synergistic utilisation of mannose and glucose by *N. frontalis* CoB3, which is dependent on the presence of glucose as primary carbon source, and strongly influenced by the concentration of mannose and the ratio to the glucose concentration.

### Tracing radio-labelled sugar metabolism confirms mannose use as co-substrate

To confirm mannose utilisation as co-substrate by *N. frontalis* CoB3, we used [^14^C]mannose to trace its metabolic fate in the presence of glucose. Given the ambiguous behaviour of arabinose in similar analyses (Fig. 3 & S3), [^14^C]arabinose was similarly evaluated. To understand the baseline metabolic flux using the radiolabelled sugars, [^14^C]glucose was included as a positive control.

Across all conditions, the majority of ^14^C was recovered in the culture medium filtrate (Fig. 5A). There was slight incorporation of the dosed ^14^C into fungal biomass from both [^14^C]glucose (∼2.4%) and [^14^C]mannose (∼2 %), but no detectable incorporation from [^14^C]arabinose. Despite the low counts, the absence of detectable ^14^C in washes 2 and 3 indicates that residual culture medium did not contribute substantially to the measured biomass-associated ^14^C.

**Figure 5.**
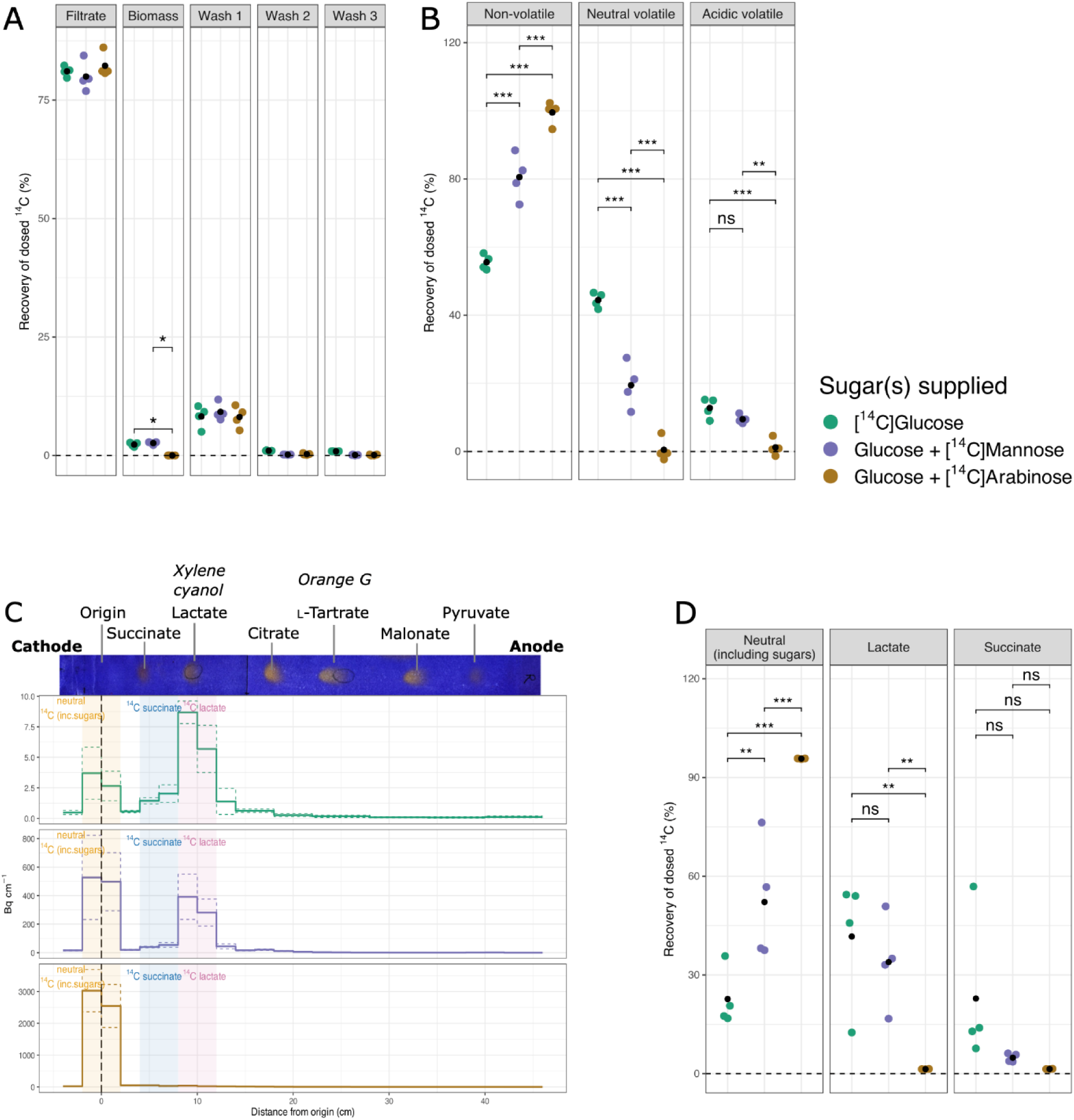
Recovery of ^14^C from dosed radiolabelled sugars after 120 hours of N. frontalis growth. Each biological replicate’s percentage recovery (coloured symbols) and the mean percentage recovery of ^14^C (black symbols) is reported for [^14^C]glucose (n=4), [^14^C]mannose (n=4), and [^14^C]arabinose (n=3). A) Fractions collected at culture harvest, B) fractions of culture filtrate. C) Mean becquerel per centimetre of Whatman paper from high voltage paper electrophoresis, with standard deviation plotted in dotted lines with corresponding colours. The origin where samples were loaded is marked at 0 cm. The identity of non-volatile ^14^C-metabolites was determined by reference to the electrophoretic mobilities of external markers stained with bromophenol blue shown here and Fig. S6, especially glucose, succinate and lactate. D)Quantified recovery of components of interest in the non-volatile fraction. Two-way ANOVA and Tukey HSD analysis was used to assess significance and p-values are given as *** p < 0.001; ** 0.001 ≤ p < 0.01; * 0.01 ≤ p < 0.05.

Further investigation of the biochemical nature of the ^14^C compounds present in the culture medium revealed distinct metabolic profiles from each radiolabelled sugar. Significantly more ^14^C was detected in the neutral volatile fraction ([^14^C]ethanol) from the [^14^C]glucose supplemented control than from either [^14^C]arabinose or [^14^C]mannose (Fig. 5B). There was no significant difference between the output of acidic volatile radioactivity ([^14^C] acetate etc.) between [^14^C]glucose and [^14^C]mannose, suggesting similar proportions of SCFAs are produced from the metabolism of these sugars. From [^14^C]glucose, a significantly lower percentage of the ^14^C was recovered in the non-volatile fraction than from [^14^C]arabinose and [^14^C]mannose (Fig. 5B). Further investigation of this fraction by high-voltage paper electrophoresis identified lactate as a major contributor of the ^14^C recovered from both [^14^C]glucose and [^14^C]mannose; surprisingly there was no significant difference in the proportion of lactate produced in these conditions. Furthermore, there was a significantly lower percentage of ^14^C present in the neutral zone from [^14^C]glucose than from [^14^C]mannose. It is likely that this is due to the amount of sugar remaining, as TLC analysis showed no glucose present whereas there was still visible mannose (Fig. S7D). Recovered ^14^C from [^14^C]arabinose is thought to be remaining unmetabolised arabinose, given its neutral nature (Fig. 5C) and the high concentration of arabinose observed to be remaining in the filtrate by TLC analysis (Fig. S7).

Together, this work provides experimental evidence of mannose utilisation hypothesised in the *Neocallimastix* metabolic model (Wilken *et al.,* 2021); however, the co-utilisation requirement indicates there are still unidentified elements integral to this metabolic pathway, such mannose transporters, receptors or regulatory elements. This highlights the fundamental gaps in knowledge of even the best-characterised AGF genus to date. These data also confirm *N. frontalis* CoB3 does not metabolise arabinose. In addition, the results indicate that there is distinct metabolic routing of carbon sources in our culture conditions with glucose compared to glucose and mannose, as judged by the measured ^14^C fermentation flux into ethanol, lactate, and SCFAs, which could be substrate-specific or linked to the amount of available metabolisable carbon.

## Discussion

AGF are integral to plant biomass degradation in the rumen, yet their strategies for sugar recognition, regulation of uptake, and metabolism remain poorly understood compared to model fungi from the Dikarya. Here, it is shown that *N. frontalis* CoB3*, C. communis* SHB and *P. edwardsiae* SHC utilise a limited repertoire of free sugars available in the rumen environment as sole carbon source, which suggests that the primary metabolism of the species assessed here is mainly glucose-and fructose driven, which could reflect their niche relative to other microbial partners in the rumen. However, utilisation of these sugars is strongly influenced by the presence of other free sugars; it was demonstrated that even sugars that are not metabolised as sole carbon source can repress fungal growth or be taken up and metabolised, when combined with glucose as carbon source. These insights in central carbon metabolism affect our understanding of the niches of AGF relative to other microbial partners in the rumen.

Of the AGF isolates assayed here, *N. frontalis* CoB3 was the most versatile and robust isolate, utilising the highest number of sugars (glucose, fructose, cellobiose, maltose, sucrose, and lactose) as the sole carbon source to support growth and was able to simultaneously remove different monosaccharides (glucose and fructose) from the culture medium (Fig. 2C,D). Interestingly, *N. frontalis* CoB3 was also the only isolate to exhibit co-metabolism of glucose and a lignocellulose-derived sugar, with a concentration-dependent synergistic utilisation of glucose and mannose, confirmed with a radiolabelled sugar tracer assay (Fig. 5). AGF are expected to produce a range of fermentation products, namely hydrogen and carbon dioxide gases, formate, acetate, ethanol, lactate, and succinate (Bauchop and Mountfort, 1981; Wilken *et al.,* 2021). Here, there was comparable percentage recovery of ^14^C in SCFAs from [^14^C]glucose- and [^14^C]mannose-dosed cultures suggesting the oxidative pathway associated with hydrogenosomal metabolism was active in both conditions (Stabler *et al.,* 2022; Schultz *et al.,* 2025). The similarly low percentages of ^14^C incorporation from both [^14^C]glucose and [^14^C]mannose into the fungal biomass is likely to be reflective of sampling at stationary growth, where growth is minimal and carbon is directed to fermentation products (Lowe, Theodorou and Trinci, 1987). Furthermore, the recovered ^14^C profile of the [^14^C]glucose control showed a substantial proportion of ^14^C recovered as ethanol and lactate, which was also observed for the [^14^C]mannose cultures (Fig. 5). This is consistent with previous observations during AGF stationary growth, their metabolism shifts to cytosolic pathways resulting in increased ethanol and lactate production as it becomes energetically less favourable to utilise the hydrogenosomal pathways which produce acetate and hydrogen gas (Stabler *et al.,* 2022; Schultz *et al.,* 2025).

Despite mannose being present at half the concentration of glucose and not being completely utilised by *N. frontalis* CoB3, similar percentages of ^14^C recovered in lactate, SCFAs, and biomass compared to [^14^C]glucose cultures (Fig. 5). This suggests that mannose is metabolised efficiently into downstream metabolic products; however, as mannose cannot be utilised as a sole carbon source by CoB3 (Fig. 1) it must function as a secondary substrate dependent on the presence of glucose. The use of mannose as a secondary substrate by *N. frontalis* CoB3 may have an ecological impact; by increasing the overall pool of lactate and ethanol it can facilitate different cross feeding interactions, supporting the growth of different bacteria and methanogens and affecting the overall fermentation efficiency of the host animal (Van Lingen *et al.,* 2017).

The inability of *N. frontalis* CoB3 to utilise mannose as a sole carbon source is contrary to the previous characterisation of *N. frontalis* RE1 strain exhibiting synergistic glucose and mannose utilisation (Stewart *et al.,* 1995). However, CoB3 is not the only strain of *N. frontalis* shown to respond to the presence of mannose despite not being able to utilise it as a sole carbon source. The strain of *N. frontalis* chemotactically characterised in the seminal work of Orpin’s group, could not utilise mannose for growth but exhibited a chemotactic response towards mannose and possessed discrete glucose- and mannose-sensing receptors (Orpin and Bauchop, 1978). Furthermore, recent genomic and transcriptomic analyses of *N. californiae* identified putative SWEET (Sugars Will Eventually be Exported Transporter) transporters predicted to mediate uptake of sugars including mannose; one was experimentally confirmed to transport mannose via heterologous expression in *S. cerevisiae,* despite *N. californiae* being unable to utilise mannose as a sole carbon source (Podolsky et al., 2021). In addition, the metabolic model of *Neocallimastix cameroonii var. lanati* predicted mannose utilisation but experimentally it was shown not to utilise mannose as the sole carbon source (Wilken *et al.,* 2021). This suggests that mannose detection could be conserved across the genus of *Neocallimastix* but the downstream transport or metabolic processing may vary at the species or strain level, potentially co-dependent on the relative concentrations of glucose and mannose.

The inhibition of *P. edwardsiae* SHC growth by mannose in the co-substrate screen of glucose and other lignocellulose-derived sugars (Fig. 3) was similar to the observed inhibition of *N. frontalis* CoB3 growth when the glucose concentration was equal to or less than that of mannose (Fig. 4) - with decreased fermentation gas production and glucose retention in the culture medium (Fig. 3E & 4C). The genus of *Piromyces* has not been characterised as extensively as *Neocallimastix,* but a similar discrepancy of this genus’s capacity of mannose utilisation is reported in the literature: *Piromyces* strain E2 was reported to utilise mannose as the sole carbon source (Teunissen *et al.,* 1992), and *Piromyces finnis* has genes for putative SWEET transporters that may be capable of mannose transport (Seppälä *et al.,* 2016) but its growth on mannose is uncertain, and *P. edwardsiae* SHC reported here cannot metabolise mannose as sole carbon source.

Therefore, it is possible that a similar mechanism and pathways of co-substrate utilisation could be present but require a higher excess of glucose (relative to mannose) than used in the initial screen performed here. Alternatively, it is possible that SHC could lack a functional metabolic pathway to catabolise mannose, and the similar structure and stereochemistry of mannose to glucose may have caused competitive inhibition of transporters and catabolic enzymes - limiting glucose catabolism and therefore, the growth of isolate SHC. However, although incomplete pathways for mannose metabolism have been reported for several AGF, e.g. *Anaeromyces robustus* (Henske *et al.,* 2018b) and *Caecomyces churrovis* (Henske *et al.,* 2017), the functional metabolic pathways are currently predicted for all AGF genomes available in the MycoCosm database (Grigoriev *et al.,* 2014), a discrepancy possibly caused by improved and updated annotations.

Surprisingly, all AGF isolates here showed replicate variability of xylose uptake in the presence of glucose (Fig 3C-E). It is possible AGF may co-import both xylose and glucose, as *N. californiae* NcSWEET1 transporter was demonstrated to facilitate transport and co-utilisation of glucose and xylose when heterologously expressed in *Saccharomyces cerevisiae,* and similar transporters were identified in the genomes of *Anaeromyces robustus,* and *P. finnis* (Podolsky *et al.,* 2021). The genes encoding enzymes for xylose catabolism via the xylose isomerase pathway have been identified in several available AGF genomes (Wilken *et al.,* 2021) and xylose isomerase has been functionally characterised in *Piromyces* sp. E2 (Harhangi *et al.,* 2003). As only *C. communis* SHB growth was significantly inhibited by the presence of xylose, this isolate could be expressing an abundance of these transporters, and the variation observed (Fig. 3) could also suggest that the expression of these SWEET transporters could be temporally regulated depending on lifecycle or environmental conditions.

Not all the lignocellulose-derived sugars tested here statistically affected the growth of *N. frontalis* CoB3, *C. communis* SHB, and *P. edwardsiae* SHC. However, from visual assessment of their growth curves, these sugars did seem to delay the AGF isolates’ transition into exponential growth (Fig. 3). It is possible that AGF may not metabolise these lignocellulose-derived sugars for growth but sense them and use them as metabolic or developmental cues, for example to trigger transitions into the next stage of their complex lifecycle similar to what has been described for the ascomycete *Aspergillus niger* (Hayer *et al.,* 2013). The required method of subculturing to maintain AGF makes it difficult to investigate this possibility, as the transferred cells are not in a synchronised growth stage. To clarify how these AGF isolates interact with lignocellulose-derived sugars which did not affect their growth by the parameters measured here, cultures could be inoculated with zoospores and ^14^C-labelled sugar used as a tracer to track the fate of the sugars at different stages of the fungal lifecycle to ascertain if they remain in the culture medium (as with arabinose; Fig. 5), attach to the fungal cells, or are metabolised and transformed into ^14^CO_2(g)_ and other metabolic products by the AGF (as with glucose and mannose; Fig. 5)(Tao *et al.,* 2016).

The preferential uptake of sugars can be used as a marker for potential niches that bacteria and fungi inhabit (Palmonari, Federiconi and Formigoni, 2024). When provided with a mixture of metabolisable sugars, the AGF isolates assayed here were found to have different sugar preferences, rates of uptake, and methods of uptake (sequential and simultaneous).

As there is limited knowledge of AGF chemotaxis in species other than *N. frontalis* (Orpin and Bountiff, 1978), we prepared the sugar mixtures with equal concentrations to minimise chemotactic bias toward a specific sugar, which could otherwise influence the fungal– substrate interaction. Each AGF species’ growth showed reduced total growth on the sugar mixtures “M1” and “M2” relative to their single sugar cultures (Fig. 1). This reduction is consistent with carbon catabolite repression (CCR), a widely conserved regulatory system in fungi, where the presence of a preferred carbon source can down-regulate pathways for alternative substrates and thereby reduce the overall metabolic efficiency. It has been demonstrated that AGF possess a CCR system (Mountfort and Asher, 1983) (Henske *et al.,* 2018a); however, the underpinning molecular mechanism has yet to be resolved and our understanding is currently based on comparison to model fungi *A. niger* or *Trichoderma reesei*, which are widely employed in biorefinery (Ries *et al.,* 2013). Notably, only *P. edwardsiae* SHC growth exhibited a clear lag phase on “M1”,. Sequential sugar utilisation in fungi such as *A. niger* can be governed by transporter kinetics rather than transcriptional repression (Mäkelä *et al.,* 2018). Therefore, if the glucose, fructose, cellobiose, and lactose sugars present in “M1” (glucose, fructose, cellobiose, and lactose) rely on overlapping transporters or sensing mechanisms – as suggested for *N. frontalis* receptors for certain sugars (Orpin and Bountiff, 1978) – competition for uptake could delay growth on less-preferred substrates. Overall, the differences in sugar utilisation hierarchies observed across *N. frontalis* CoB3, *C. communis* SHB, and *P. edwardsiae* SHC suggest variation in their carbon-source regulatory networks, which may include differences in CCR strength, transporter repertoires, or sensing mechanisms.

The AGF isolates here were maintained in a nutrient-rich medium with wheat straw as the carbon source. To investigate whether a substrate ‘priming’ effect influences AGF sugar utilisation – a process previously observed in the rumen bacteria *Bacteroides* whereby growth on complex polysaccharides accelerated subsequent glycan breakdown and product uptake (Klassen *et al.,* 2021) – AGF isolate growth was screened on the selection of soluble sugars from inocula pre-grown on glucose (a simple sugar) or wheat straw (a complex lignocellulose). From the phenotypic data collected here, the carbon source of the inoculum did not seem to influence the sugar utilisation of the subsequent culture. However, it did affect the subsequent culture’s performance in terms of total accumulated fermentation gas pressure. The accumulated gas pressure recorded for cultures inoculated from the simple sugar source (glucose) produced significantly less fermentation gas than the inoculum from complex lignocellulose (wheat straw) for subsequent growth on glucose (all isolates), cellobiose (SHB and SHC), and maltose (SHC). This reduction in fermentation gas could indicate that the fungal isolates became more efficient in glucose utilisation (leading to a lower gas generation to fungal biomass ratio, given that gas pressure is an indirect proxy for growth). Alternatively, as the glucose-pregrown AGF cultures grew faster (Fig. 1), when used as an inoculum at 3 days old, these cultures could have fewer zoospores than the cultures pregrown on wheat straw, resulting in less effective growth.

In summary, this work has shown that AGF isolates *N. frontalis* CoB3, *C. communis* SHB, and *P. edwardsiae* SHC differ in fundamental aspects of simple sugar metabolism, including clear differences in substrate preference and the presence of novel co-substrate dependencies. These findings provide a useful first framework for considering each isolate’s native potential as an inoculant for lignocellulosic waste valorisation in anaerobic digestors. It also offers a basis for a broader understanding of the ecological roles of AGF in the complex rumen microbiome environment, required to create pathways towards building simplified microbial communities to explore rumen cross-feeding interactions and reducing ruminant-farming associated methane emissions. Further work will be required to resolve the mechanisms underpinning these co-substrate dependencies and to understand how processes such as CCR and chemotaxis may influence AGF performance when confronted with complex, recalcitrant substrates.

## CRediT authorship contribution statement

**JLM:** Writing – review & editing, Writing – original draft, Investigation, Visualization, Validation, Methodology, Formal analysis, Data curation, Conceptualization, Funding acquisition. **HH:** Investigation, Formal analysis, Data curation, **SCF:** Writing – review & editing, Conceptualization, Methodology, Supervision, Project administration, Funding acquisition. **JMvM:** Writing – review & editing, Conceptualization, Methodology, Formal analysis, Data curation, Supervision, Project administration, Funding acquisition.

## Acknowledgements

The authors thank the Royal Society for support via URF\R1\231686, and the UKRI Biotechnology and Biological Sciences Research Council (BBSRC) through the EASTBIO DTP, grant number BB/T00875X/1.

## References

Akin, D.E. and Borneman, W.S. (1990) ‘Role of rumen fungi in fiber degradation’, Journal of Dairy Science, 73(10), pp. 3023–3032.

Bauchop, T., 1979. Rumen anaerobic fungi of cattle and sheep. Applied and Environmental Microbiology, 38(1), pp.148–158.

Bauchop, T. and Mountfort, D.O., 1981. Cellulose fermentation by a rumen anaerobic fungus in both the absence and the presence of rumen methanogens. Applied and environmental microbiology, 42(6), pp.1103–1110.

Braune, R. (1913) ‘Untersuchungen über die im Wiederkäuermagen vorkommenden Protozoen’, Archiv für Protistenkunde, 32, pp. 111–170.

Fliegerova, K.O., Podmirseg, S.M., Vinzelj, J., Grilli, D.J., Kvasnová, S., Schierová, D., Sechovcová, H., Mrázek, J., Siddi, G., Arenas, G.N. and Moniello, G., 2021. The effect of a high-grain diet on the rumen microbiome of goats with a special focus on anaerobic fungi. Microorganisms, 9(1), p.157.

Franková, L. and Fry, S.C., 2021. Hemicellulose-remodelling transglycanase activities from charophytes: towards the evolution of the land-plant cell wall. The Plant Journal, 108(1), pp.7–28.

Fry, S.C., 1988. The growing plant cell wall: chemical and metabolic analysis (pp. 333-pp).

Fry, S.C., 2020. High-voltage paper electrophoresis (HVPE). In The Plant Cell Wall: Methods and Protocols (pp. 1–31). New York, NY: Springer New York.

Grenet, E. and Barry, P., 1988. Colonization of thick-walled plant tissues by anaerobic fungi. Animal feed science and technology, 19(1-2), pp.25–31.

Gruninger, R.J., Nguyen, T.T., Reid, I.D., Yanke, J.L., Wang, P., Abbott, D.W., Tsang, A. and McAllister, T., 2018. Application of transcriptomics to compare the carbohydrate active enzymes that are expressed by diverse genera of anaerobic fungi to degrade plant cell wall carbohydrates. Frontiers in microbiology, 9, p.1581.

Haitjema, C.H., Gilmore, S.P., Henske, J.K., Solomon, K.V., De Groot, R., Kuo, A., Mondo, S.J., Salamov, A.A., LaButti, K., Zhao, Z. and Chiniquy, J., 2017. A parts list for fungal cellulosomes revealed by comparative genomics. Nature microbiology, 2(8), pp.1–8.

Han, X., Li, B., Wang, X., Chen, Y. and Yang, Y., 2019. Effect of dietary concentrate to forage ratios on ruminal bacterial and anaerobic fungal populations of cashmere goats. Anaerobe, 59, pp.118–125.

Hanafy, R.A., Lanjekar, V.B., Dhakephalkar, P.K., Callaghan, T.M., Dagar, S.S., Griffith, G.W., Elshahed, M.S. and Youssef, N.H., 2020. Seven new Neocallimastigomycota genera from wild, zoo-housed, and domesticated herbivores greatly expand the taxonomic diversity of the phylum. Mycologia, 112(6), pp.1212–1239.

Harhangi, H.R., Akhmanova, A.S., Emmens, R., van der Drift, C., de Laat, W.T., van Dijken, J.P., Jetten, M.S., Pronk, J.T. and Op den Camp, H.J., 2003. Xylose metabolism in the anaerobic fungus Piromyces sp. strain E2 follows the bacterial pathway. Archives of microbiology, 180(2), pp.134–141.

Hayer, K., Stratford, M. and Archer, D.B., 2013. Structural features of sugars that trigger or support conidial germination in the filamentous fungus Aspergillus niger. Applied and environmental microbiology, 79(22), pp.6924–6931.

Henske, J.K., Gilmore, S.P., Haitjema, C.H., Solomon, K.V. and O’Malley, M.A., 2018. Biomass-degrading enzymes are catabolite repressed in anaerobic gut fungi. AIChE Journal, 64(12), pp.4263–4270.

Henske, J.K., Gilmore, S.P., Knop, D., Cunningham, F.J., Sexton, J.A., Smallwood, C.R., Shutthanandan, V., Evans, J.E., Theodorou, M.K. and O’Malley, M.A., 2017. Transcriptomic characterization of Caecomyces churrovis: a novel, non-rhizoid-forming lignocellulolytic anaerobic fungus. Biotechnology for Biofuels, 10(1), p.305.

Henske, J.K., Wilken, S.E., Solomon, K.V., Smallwood, C.R., Shutthanandan, V., Evans, J.E., Theodorou, M.K. and O’Malley, M.A., 2018. Metabolic characterization of anaerobic fungi provides a path forward for bioprocessing of crude lignocellulose. Biotechnology and Bioengineering, 115(4), pp.874–884.

Hess, M., Paul, S.S., Puniya, A.K., Van der Giezen, M., Shaw, C., Edwards, J.E. and Fliegerová, K., 2020. Anaerobic fungi: past, present, and future. Frontiers in Microbiology, 11, p.584893.

Grigoriev, I.V., Nikitin, R., Haridas, S., Kuo, A., Ohm, R., Otillar, R., Riley, R., Salamov, A., Zhao, X., Korzeniewski, F. and Smirnova, T., 2014. MycoCosm portal: gearing up for 1000 fungal genomes. Nucleic acids research, 42(D1), pp.D699–D704.

Joblin, K.N. and Naylor, G.E., 1993. Inhibition of the rumen anaerobic fungus Neocallimastix frontalis by fermentation products. Letters in applied microbiology, 16(5), pp.254–256.

Jones, A.L., Clayborn, J., Pribil, E., Foote, A.P., Montogomery, D., Elshahed, M.S. and Youssef, N.H., 2023. Temporal progression of anaerobic fungal communities in dairy calves from birth to maturity. Environmental Microbiology, 25(11), pp.2088–2101.

Klassen, L., Reintjes, G., Tingley, J.P., Jones, D.R., Hehemann, J.H., Smith, A.D., Schwinghamer, T.D., Arnosti, C., Jin, L., Alexander, T.W. and Amundsen, C., 2021. Quantifying fluorescent glycan uptake to elucidate strain-level variability in foraging behaviors of rumen bacteria. Microbiome, 9(1), p.23.

Lankiewicz, T.S., Choudhary, H., Gao, Y., Amer, B., Lillington, S.P., Leggieri, P.A., Brown, J.L., Swift, C.L., Lipzen, A., Na, H. and Amirebrahimi, M., 2023. Lignin deconstruction by anaerobic fungi. Nature Microbiology, 8(4), pp.596–610.

Lillington, S.P., Chrisler, W., Haitjema, C.H., Gilmore, S.P., Smallwood, C.R., Shutthanandan, V., Evans, J.E. and O’Malley, M.A., 2021. Cellulosome localization patterns vary across life stages of anaerobic fungi. Mbio, 12(3), pp.10–1128.

Lowe, S.E., Theodorou, M.K. and Trinci, A.P., 1987. Growth and fermentation of an anaerobic rumen fungus on various carbon sources and effect of temperature on development. Applied and environmental microbiology, 53(6), pp.1210–1215.

Mäkelä, M.R., Aguilar-Pontes, M.V., van Rossen-Uffink, D., Peng, M. and de Vries, R.P., 2018. The fungus Aspergillus niger consumes sugars in a sequential manner that is not mediated by the carbon catabolite repressor CreA. Scientific reports, 8(1), p.6655.

Meili, C.H., TagElDein, M.A., Jones, A.L., Moon, C.D., Andrews, C., Kirk, M.R., Janssen, P.H., J. Yeoman, C., Grace, S., Borgogna, J.L.C. and Foote, A.P., 2024. Diversity and community structure of anaerobic gut fungi in the rumen of wild and domesticated herbivores. Applied and Environmental Microbiology, 90(2), pp.e01492–23.

Mountfort, D.O. and Asher, R.A., 1983. Role of catabolite regulatory mechanisms in control of carbohydrate utilization by the rumen anaerobic fungus Neocallimastix frontalis. Applied and environmental microbiology, 46(6), pp.1331–1338.

Nagaraja, T.G. (2016) ‘Microbiology of the rumen’, in Rumenology. Springer, pp. 39–61.

Orpin, C.G., 1975. Studies on the rumen flagellate Neocallimastix frontalis. Microbiology, 91(2), pp.249–262.

Orpin, C.G. and Bountiff, L., 1978. Zoospore chemotaxis in the rumen phycomycete Neocallimastix frontalis. Microbiology, 104(1), pp.113–122.

Palmonari, A., Federiconi, A. and Formigoni, A., 2024. Animal board invited review: The effect of diet on rumen microbial composition in dairy cows. animal, 18(10), p.101319.

Podolsky, I.A., Seppälä, S., Xu, H., Jin, Y.S. and O’Malley, M.A., 2021. A SWEET surprise: anaerobic fungal sugar transporters and chimeras enhance sugar uptake in yeast. Metabolic Engineering, 66, pp.137–147.

Ries, L., Pullan, S.T., Delmas, S., Malla, S., Blythe, M.J. and Archer, D.B., 2013. Genome-wide transcriptional response of Trichoderma reesei to lignocellulose using RNA sequencing and comparison with Aspergillus niger. BMC genomics, 14(1), p.541.

Seppälä, S., Knop, D., Solomon, K.V. and O’Malley, M.A., 2017. The importance of sourcing enzymes from non-conventional fungi for metabolic engineering and biomass breakdown. Metabolic engineering, 44, pp.45–59.

Seppälä, S., Solomon, K.V., Gilmore, S.P., Henske, J.K. and O’Malley, M.A., 2016. Mapping the membrane proteome of anaerobic gut fungi identifies a wealth of carbohydrate binding proteins and transporters. Microbial cell factories, 15(1), p.212.

Schulz, K.E., Scholz, D., Sikirić, A.R., Rambow, D., Neumann, A. and Ochsenreither, K., 2025. Anaerobic gut fungi as biocatalysts: metabolic and physiological analysis of anaerobic gut fungi under diverse cultivation conditions. Frontiers in Microbiology, 16, p.1662047.

Shen, S., Matthews, J.L., Li, S. and van Munster, J.M., 2026. Anaerobic fungi Caecomyces communis, Neocallimastix frontalis and Piromyces edwardsiae sp. nov. have distinct effects on plant fibres during digestion. Royal Society Open Science, 13(3).

Sluiter, J.B., Ruiz, R.O., Scarlata, C.J., Sluiter, A.D. and Templeton, D.W., 2010. Compositional analysis of lignocellulosic feedstocks. 1. Review and description of methods. Journal of agricultural and food chemistry, 58(16), pp.9043–9053.

Solomon, K.V., Haitjema, C.H., Henske, J.K., Gilmore, S.P., Borges-Rivera, D., Lipzen, A., Brewer, H.M., Purvine, S.O., Wright, A.T., Theodorou, M.K. and Grigoriev, I.V., 2016. Early-branching gut fungi possess a large, comprehensive array of biomass-degrading enzymes. Science, 351(6278), pp.1192–1195.

Stabel, M., Haack, K., Lübbert, H., Greif, M., Gorenflo, P., Aliyu, H. and Ochsenreither, K., 2022. Metabolic shift towards increased biohydrogen production during dark fermentation in the anaerobic fungus Neocallimastix cameroonii G341. Biotechnology for biofuels and bioproducts, 15(1), p.96.

Stewart, C.S., Duncan, S.H., Richardson, A.J., Calder, A.G. and Dewey, P.J., 1995. The effect of the presence of glucose on the fermentation of mannose by the anaerobic fungus Neocallimastix frontalis strain RE1. FEMS microbiology letters, 127(1-2), pp.57–63.

Tamminga, S. and Van Vuuren, A.M., 1988. Formation and utilization of end products of lignocellulose degradation in ruminants. Animal Feed Science and Technology, 21(2-4), pp.141–159.

Teunissen, M.J., de Kort, G.V., Op den Camp, H.J. and Huis in’t Veld, J.H., 1992. Production of cellulolytic and xylanolytic enzymes during growth of the anaerobic fungus Piromyces sp. on different substrates. Microbiology, 138(8), pp.1657–1664.

Theodorou, M.K., Brookman, J. and Trinci, A.P., 2005. Anaerobic fungi. In Methods in gut microbial ecology for ruminants (pp. 55–66). Dordrecht: Springer Netherlands.

Theodorou, M.K., Davies, D.R., Nielsen, B.B., Lawrence, M.I. and Trinci, A.P., 1995. Determination of growth of anaerobic fungi on soluble and cellulosic substrates using a pressure transducer. Microbiology, 141(3), pp.671–678.

Theodorou, M.K., Mennim, G., Davies, D.R., Zhu, W.Y., Trinci, A.P. and Brookman, J.L., 1996. Anaerobic fungi in the digestive tract of mammalian herbivores and their potential for exploitation. Proceedings of the Nutrition Society, 55(3), pp.913–926

Tao, J., Diaz, R.K., Teixeira, C.R. and Hackmann, T.J., 2016. Transport of a fluorescent analogue of glucose (2-NBDG) versus radiolabeled sugars by rumen bacteria and Escherichia coli. Biochemistry, 55(18), pp.2578–2589.

Van Lingen, H.J., Edwards, J.E., Vaidya, J.D., Van Gastelen, S., Saccenti, E., Van den Bogert, B., Bannink, A., Smidt, H., Plugge, C.M. and Dijkstra, J., 2017. Diurnal dynamics of gaseous and dissolved metabolites and microbiota composition in the bovine rumen. Frontiers in microbiology, 8, p.425.

Wilken, S.E., Leggieri, P.A., Kerdman-Andrade, C., Reilly, M., Theodorou, M.K. and O’Malley, M.A., 2020. An Arduino based automatic pressure evaluation system to quantify growth of non-model anaerobes in culture. AIChE Journal, 66(12), p.e16540.

Wilken, S.E., Monk, J.M., Leggieri, P.A., Lawson, C.E., Lankiewicz, T.S., Seppälä, S., Daum, C.G., Jenkins, J., Lipzen, A.M., Mondo, S.J. and Barry, K.W., 2021. Experimentally validated reconstruction and analysis of a genome-scale metabolic model of an anaerobic Neocallimastigomycota fungus. Msystems, 6(1), pp.10–1128.

Youssef, N.H., Couger, M.B., Struchtemeyer, C.G., Liggenstoffer, A.S., Prade, R.A., Najar, F.Z., Atiyeh, H.K., Wilkins, M.R. and Elshahed, M.S., 2013. The genome of the anaerobic fungus Orpinomyces sp. strain C1A reveals the unique evolutionary history of a remarkable plant biomass degrader. Applied and environmental microbiology, 79(15), pp.4620–4634

