## Supplementary material for "Comparative sugar utilisation and metabolism of mannose as co-substrate indicate flexibility in carbon metabolism in anaerobic gut fungi"

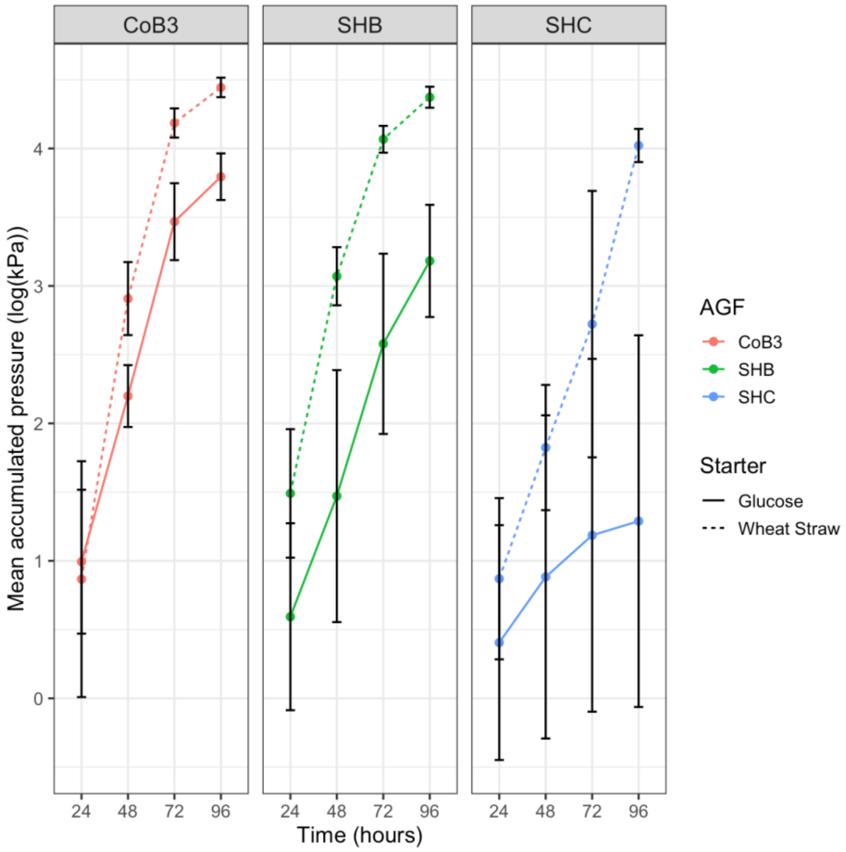


***Supplementary Figure 1. Example of $log_{10}$ transformation of mean accumulated gas pressure of AGF isolates used to identify exponential growth stage.*** Exponential growth rates were identified using the $log_{10}$ transformed mean accumulated gas pressure of AGF isolates and visually identifying the region of linearity. To this region, linear regression was then applied, and the resulting slope was used to estimate the exponential growth rate.

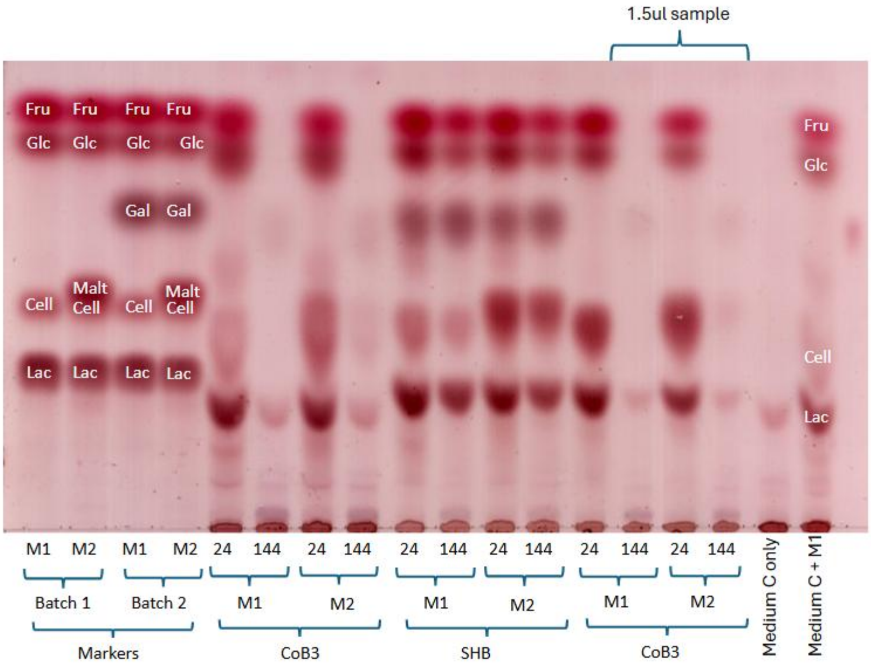


***Supplementary Figure 2. thin-layer chromatography validation of retention factors of sugars dissolved in water versus culture medium.***

To confirm that the correct disaccharides were added to the markers, two independent batches of “M1” and “M2” sugar marker mixtures were prepared. Galactose, the presumed product of lactose hydrolysis, was added to “Batch 2” to confirm its identity.

Following the protocol described in the materials and methods, 3 µL of the sample was loaded for a different biological replicate of CoB3 and SHB used in Figure 3 to identify if it was sample-specific. As the retention factors ($R_F$s) were still differing between samples, 1.5 µL of CoB3 culture medium was added to see if this was due to overloading the sample on the plate, and consequently, if the lower sugar concentration loaded was still visible on the plate. To confirm that it was the culture medium that was affecting the mobility of sugars and not fungal modification, 3 µL medium C and “M1” (1:1 (v/v)) and 3 µL of medium C with no sugar and fungi were loaded onto the plate. This medium was of the same batch used to grow the AGF isolates. Blank medium C had a faint band; however, owing to the undefined nature of the rumen fluid, this could not be identified. The repeated differing retention factors of sugars dissolved in water compared to culture medium, and the perturbation of retention of sugars “M1” in the medium, confirmed the correct identity assignment of the sugars in experimental culture conditions (Fig. 2).

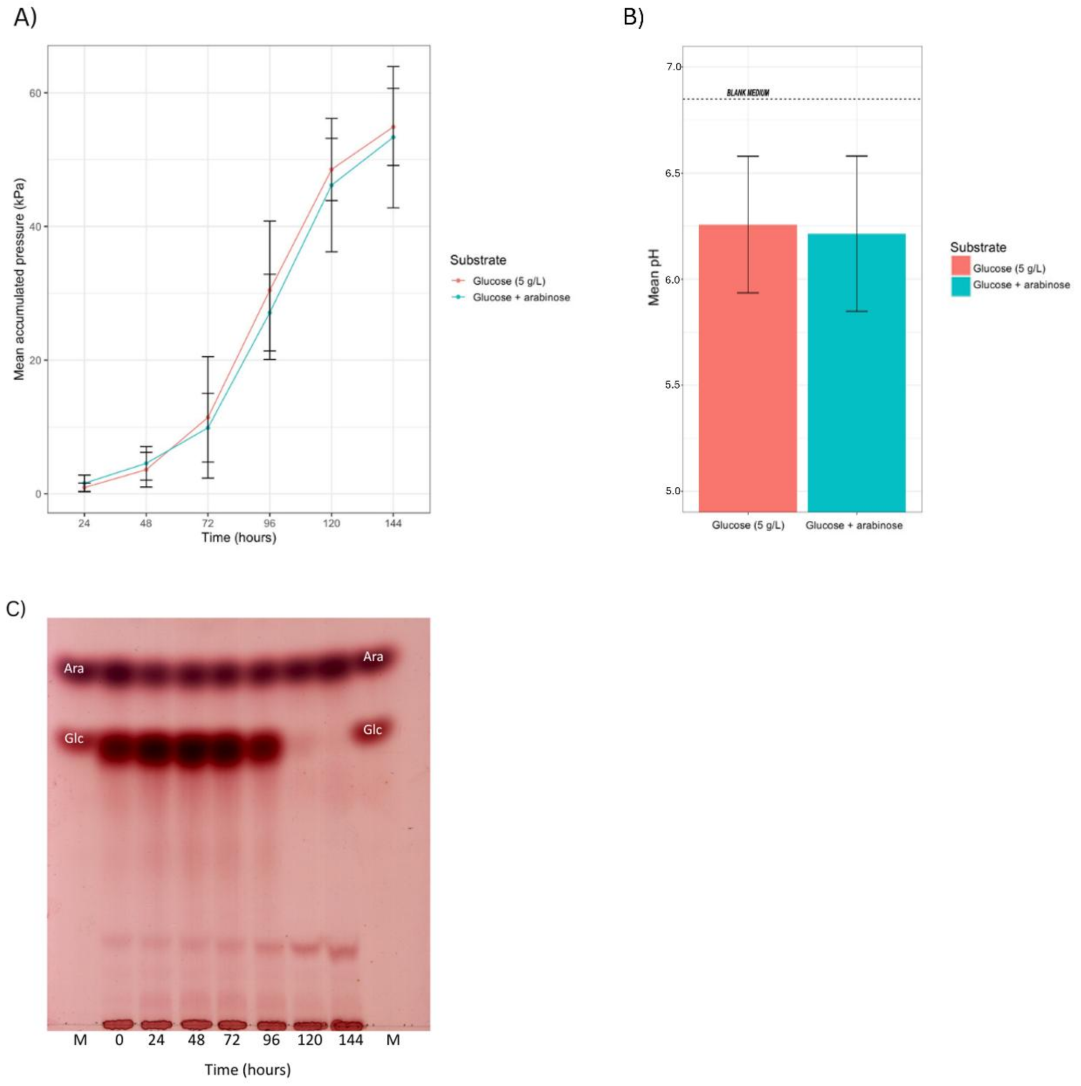


***Supplementary Figure 3. Growth of fungal isolate N. frontalis CoB3 in co-substrate culture of glucose and arabinose.*** *Glucose concentration was 5 g $L^{-1}$ and arabinose concentration was 2.5 g $L^{-1}$. Values reported are mean ± stdev (n=5). A) comparison of fermentation gas accumulation of glucose and arabinose to glucose only substrate conditions and B) pH of culture supernatant after 144 h of growth. C) Thin-layer chromatography of culture medium sampled every 24 hours of a representative replicate. '0 h' is a sample of the culture medium supplemented with the sugar co-substrates that was sampled before fungal inoculation and pooled.*

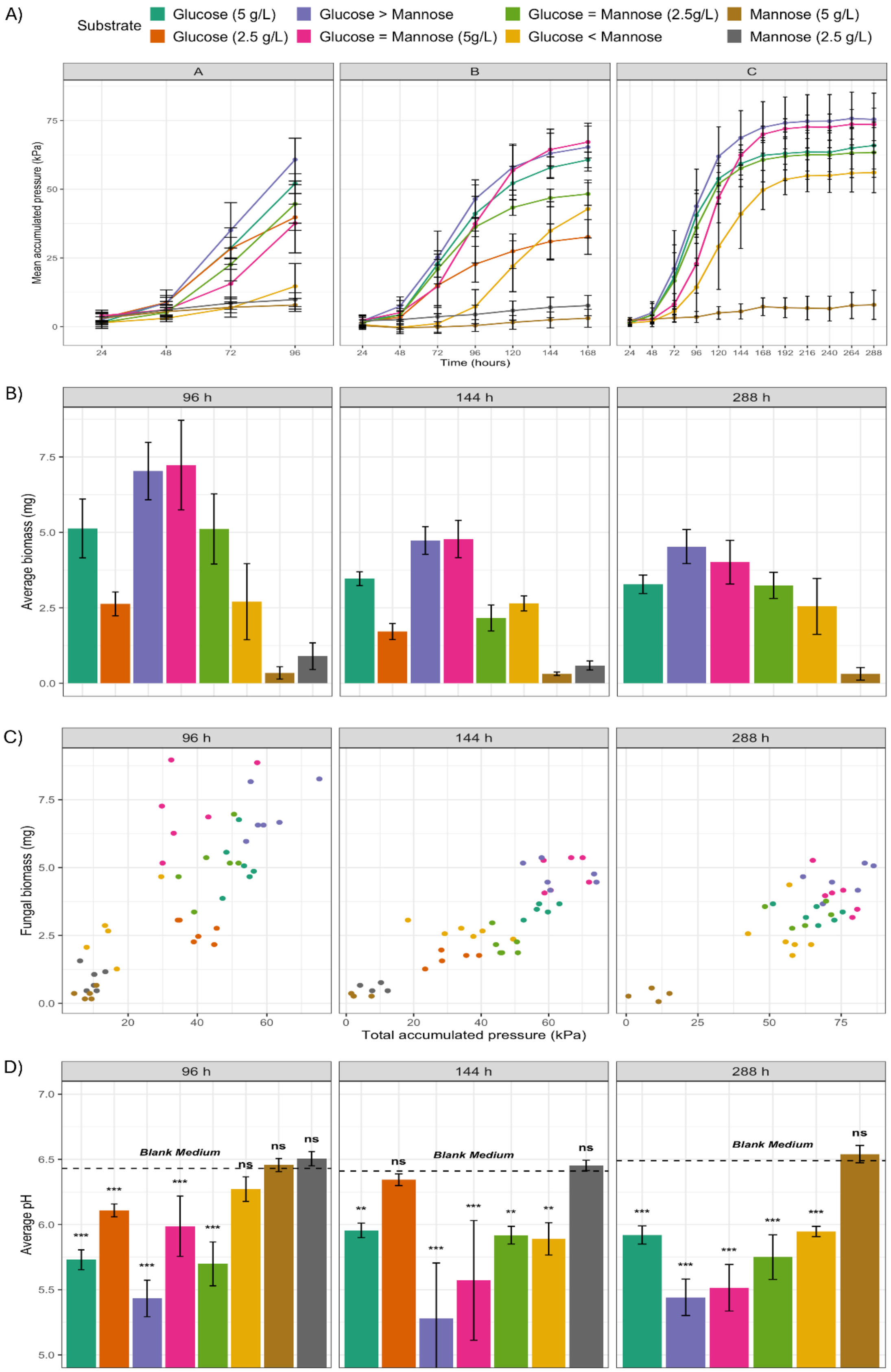


***Supplementary Figure 4. N. frontalis biomass during growth on glucose and mannose correlates with gas production.*** *A) Fermentation gas accumulation of cultures terminated at 96, 144 and 288h; B) Fungal biomass (dry weight) upon termination C) comparison of production of fermentation gas with biomass D) pH of culture supernatant. Values reported are mean ± stdev (n=5). For all statistical tests p-values given as *** $p < 0.001$; ** $0.001 \le p < 0.01$; * $0.01 \le p < 0.05$. The concentrations of sugars in cultures were: glucose (5 g $L^{-1}$) > mannose (2.5 g $L^{-1}$) and glucose (2.5 g $L^{-1}$) < mannose (5 g $L^{-1}$).*

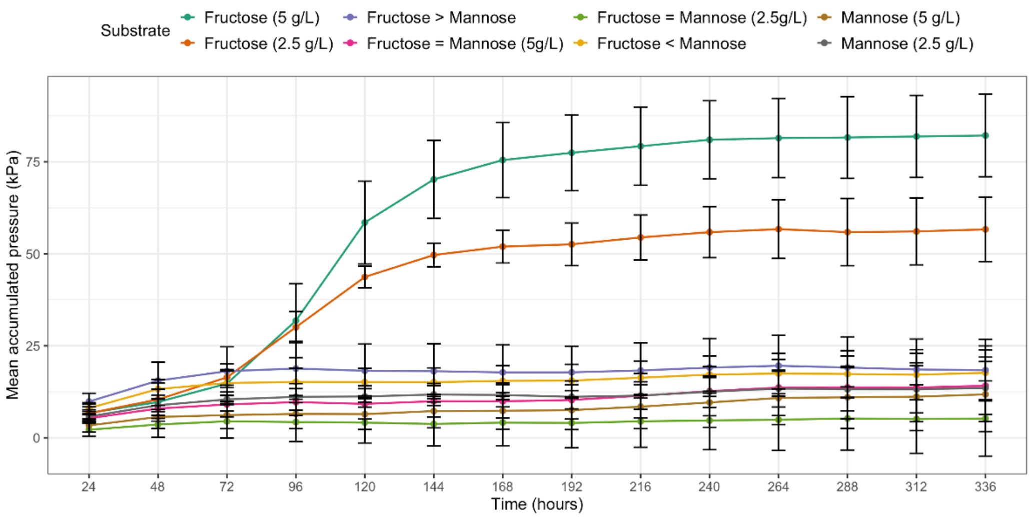


***Supplementary Figure 5. Fermentation gas production during N. frontalis CoB3 growth on fructose and mannose.*** *Fermentation gas accumulation of cultures is given as mean ± stdev (n=5). The concentrations of sugars in cultures were: fructose (5 g $L^{-1}$) > mannose (2.5 g $L^{-1}$) and fructose (2.5 g $L^{-1}$) < mannose (5 g $L^{-1}$).*

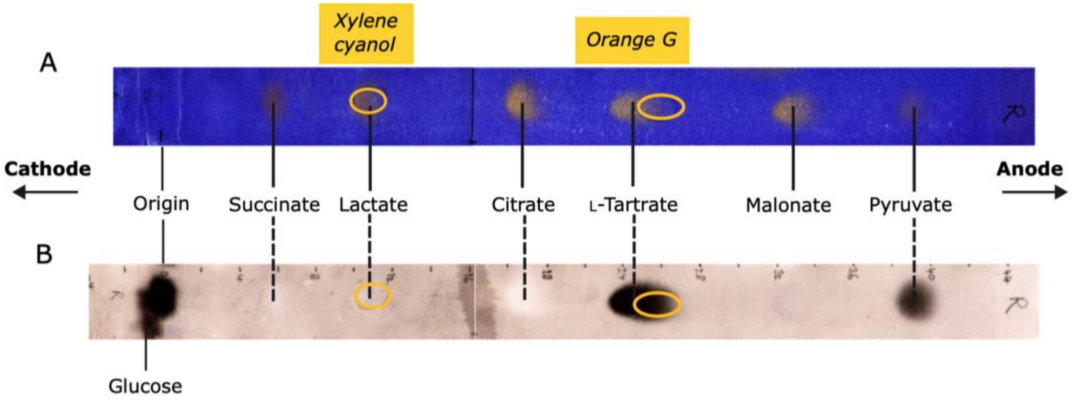


***Supplementary Figure 6. External markers used to aid in identification of 14C-metabolites recovered from non-volatile fraction.*** *Marker tracks similar to this were run alongside each radioactive track. Shown are two images of the same external marker strip stained with A) bromophenol blue and B) silver nitrate. The positions of the coloured markers (xylene cyanol and orange G; identified here with orange circles and labels), which were included in all tracks, were pencilled on the paper prior to staining. The colourless markers (glucose plus the named anions), as visualised by staining, are labelled in black. Some of the anions (indicated with -----) produced a white spot visible against the background on staining with silver nitrate.*

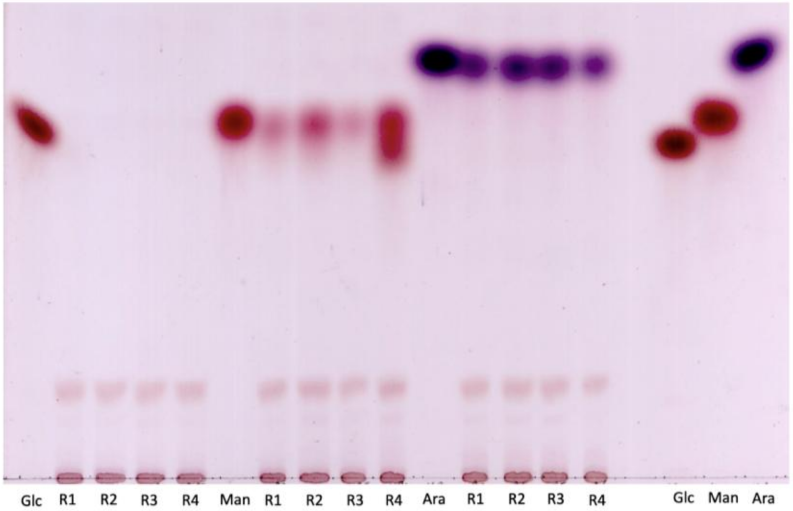


***Supplementary Figure 7. Thin layer chromatography of N. frontalis culture medium filtrate dosed with radiolabelled sugars after 120 hours of growth.*** *Each harvested culture was assayed to identify if there were still measurable glucose and mannose/arabinose present (if supplemented). Single lane markers of 2.5 µg of glucose, mannose, and arabinose were loaded for reference.*
